# *TP53* abnormalities correlate with immune infiltration and are associated with response to flotetuzumab, an investigational immunotherapy, in acute myeloid leukemia

**DOI:** 10.1101/2020.02.28.961391

**Authors:** Catherine Lai, Jayakumar Vadakekolathu, Stephen Reeder, Sarah E. Church, Tressa Hood, Ibrahim Aldoss, John Godwin, Matthew J. Wieduwilt, Martha Arellano, John Muth, Farhad Ravandi, Kendra Sweet, Heidi Altmann, Gemma A. Foulds, Friedrich Stölzel, Jan Moritz Middeke, Marilena Ciciarello, Antonio Curti, Peter J.M. Valk, Bob Löwenberg, Martin Bornhäuser, John F. DiPersio, Jan K. Davidson-Moncada, Sergio Rutella

## Abstract

**Purpose:** Somatic *TP53* mutations and 17p deletions with genomic loss of *TP53* occur in 37-46% of acute myeloid leukemia (AML) cases with adverse risk cytogenetics and are associated with primary induction failure (PIF), high risk of relapse and dismal prognosis. Herein, we aimed to characterize the immune landscape of *TP53* mutated AML and to determine whether *TP53* abnormalities identify a patient subgroup that may benefit from T-cell targeting immunotherapy approaches.

**Experimental Design:** The NanoString Pan-Cancer IO 360™ assay was used for the immune transcriptomic analysis of 64 diagnostic bone marrow (BM) samples from adults with *TP53* mutated AML (n=42) or *TP53* wild type AML (n=22), and 35 BM samples from heavily pretreated patients with relapsed/refractory (R/R) AML (11 cases with *TP53* mutations and/or 17p deletion with genomic loss of *TP53*) who received immunotherapy with flotetuzumab, an investigational CD123×CD3 bispecific DART^®^ molecule (NCT02152956). In silico data series included The Cancer Genome Atlas (TCGA) cohort and a Dutch–Belgian Cooperative Trial Group for Hematology–Oncology (HOVON) cohort.

**Results:** All TCGA cases with *TP53* mutations (n=13) expressed higher levels of negative immune checkpoints, inflammatory chemokines, interferon (IFN)-γ-inducible molecules, and had a higher tumor inflammation signature (TIS) score, compared with TCGA cases with other risk-defining molecular lesions. The comparison between *TP53* mutated and *TP53* wild type primary BM samples showed higher expression of *IFNG, FoxP3*, immune checkpoints and markers of exhaustion and senescence in the former cohort and allowed the computation of a 34-gene immune classifier prognostic for overall survival. *In vitro* modeling experiments with AML cell lines showed heightened expression of IFN-γ and inflammation pathway genes in KG-1 cells (loss-of-function mutation of *TP53*) compared with Kasumi-1 cells (gain-of-function mutation of *TP53*). Finally, 5 out of 11 (45.5%) patients with R/R AML and *TP53* abnormalities showed evidence of anti-leukemic activity of flotetuzumab immunotherapy and had higher TIS, *FoxP3*, CD8 T-cell abundance, inflammatory chemokine and *PD1* gene expression scores at baseline compared with non-responders.

**Conclusions:** This study provides evidence for a correlation between IFN-γ-dominant immune subtypes and *TP53* abnormalities. The anti-leukemic activity with flotetuzumab encourages further study of this immunotherapeutic approach in this patient subgroup.

## Introduction

Acute myeloid leukemia (AML) is a molecularly and clinically heterogeneous disease. Remission rates in newly diagnosed patients are modest and approximately 50% of patients relapse following remission. The patients with the worst outcomes are those with refractory disease, including primary induction failure (PIF) patients that fail more than one induction attempt (1). Somatic *TP53* mutations and deletions of 17p, to which *TP53* is mapped, occur in 5-10% of *de novo* AML cases (2-4) and in up to 37-46% of patients with adverse-risk cytogenetics and treatment-related myeloid neoplasms (5-7). Newly diagnosed *TP53* mutated patients have response rates to cytarabine-based chemotherapy combinations between 14-42% with a median overall survival (OS) of 2-12 months (2,6,8). Patients with 17p (*TP53*) deletion have a median OS time of 5 months and a 2-year disease-free survival (DFS) and 2-year OS time of 0% (4). While newer studies using a backbone of a hypomethylating agent or low dose cytarabine in combination with venetoclax have shown complete remission (CR) rates of 47% and 30%, respectively, in newly diagnosed *TP53* mutated AML patients, these CR rates and the median OS (7.2 and 3.7 months) are still inferior compared to the remaining cohort of patients (9,10). In the relapsed and primary refractory population, *TP53* mutations are highly enriched and response rates to current standard of care are even lower at approximately 20% with standard salvage cytotoxic regimens (6,11-13). Moreover, many patients with mutated *TP53* and/or 17p deletion have higher age and/or reduced performance status and therefore only few of them are candidates for allogeneic hematopoietic stem cell transplantation (HSCT), which offers the highest curative potential (14).

Emerging evidence implicates mutant *TP53* in activating genes involved in immune responses and inflammation (15). Studies in mice have shown that *TP53* inactivation in murine T cells augments differentiation to T helper type (Th)17 cells, thereby promoting spontaneous autoimmunity (16), and in contrast, active *TP53* can suppress inflammatory responses through the inhibition of tumor necrosis factor (TNF) transcription (17). Cancer-specific loss of *TP53* expression in lung and pancreas tumor models protects from immune-mediated elimination through the recruitment of both myeloid cells and regulatory T (Treg) cells (18). In human tumors, *TP53* mutations are enriched in the immune favorable, Th1-dominant phenotype of breast cancer, which expresses high levels of negative immune checkpoints programmed death receptor ligand 1 (PD-L1) and programmed death receptor 1 (PD1), as well as immune suppressive mediators such as indoleamine 2,3-dioxygenase-1 (19). Accumulation of *TP53* in lung tumor cells has been correlated with increased PD-L1 expression and with poor recurrence-free survival and overall survival (OS) (20). Similarly, *TP53* mutations are associated with persistent STAT3 signaling, increased cancer infiltration with GATA3^+^ Th2 cells and shorter OS in patients with pancreatic adenocarcinoma (21). Intriguingly, higher proportions of PD-L1-expressing CD8^+^ T cells, higher tumor mutational burden (TMB) and increased expression of T cell effector genes and interferon (IFN)-γ–related genes have been associated with favorable responses to pembrolizumab immunotherapy in patients with p53 mutated lung cancer (22). We have recently identified microenvironmental immune gene sets that capture elements of IFN-γ-driven biology and stratify newly diagnosed AML into an immune-infiltrated and an immune-depleted subtype (23). Our immune classifier increased the accuracy of survival prediction in patients receiving chemotherapy beyond the current capabilities of individual molecular markers. Herein, we aimed to investigate whether p53 mutations shape the immune landscape of AML and whether they identify patients that derive benefit from T cell-targeting immunotherapy approaches.

## Patients and Methods

### Patients’ demographics and study approval

Patient and disease characteristics as well as induction treatment regimens are summarized in **Table 1**. *TP53* mutational status is shown in **Supplemental Tables 1-3**. The first cohort consisted of 40 primary bone marrow (BM) samples from patients with newly diagnosed, *TP53* mutated AML treated with curative intent (SAL cohort). The second cohort included 24 primary BM samples from patients with newly diagnosed AML treated with curative intent (Bologna cohort; 2 cases with mutated *TP53*). The third cohort consisted of 35 primary BM samples collected from 27 patients with PIF or early relapse AML (CR1 < 6 months) and from 8 patients with late relapsed AML (CR1 ≥6 months) treated with flotetuzumab, an investigational CD123×CD3 bispecific DART^®^ molecule, at the recommended phase 2 dose (500 ng/kg/day) on the CP-MGD006-01 clinical trial (NCT#02152956). Patients were ineligible to receive flotetuzumab if they had been treated with a prior HSCT. Eleven patients from the flotetuzumab cohort (9 PIF/early relapse and 2 late relapse) harbored *TP53* mutations or 17p deletions with genomic loss of *TP53*. Patients received a lead-in dose of flotetuzumab during week (W) 1, followed by 500 ng/kg/day during weeks 2-4 of cycle 1, and a 4-day on/3-day off schedule for cycle 2 and beyond. Disease status was assessed by modified IWG criteria. However, it is presently unknown whether current criteria for clinical response in acute leukemia, including the proper timing of response evaluation, are adequate to define response to immunotherapies with bispecific antibodies (24). In the present study, anti-leukemic activity (ALA) of flotetuzumab was used as a surrogate study endpoint that might reflect disease control rates. ALA was defined as either CR, CR with partial hematological recovery (CRh), CR with incomplete hematological recovery (CRi), morphological leukemia-free state (MLFS) or other benefit (OB; >30% reduction of BM blasts from baseline) at the end of cycle 1. Human studies were approved by the Institutional Review Board (IRB) at the Study Alliance Leukemia (Germany) and the University of Bologna (Italy), and by the IRBs of the Institutions participating to the flotetuzumab immunotherapy clinical trial. Written informed consent was received from all participants prior to inclusion in the study.

**Table 1:** Patient series.

|  | **SAL** | **Bologna** | **TCGA<sup>^</sup>** | **HOVON** |
| --- | --- | --- | --- | --- |
|  | **Wet lab** |  | **In silico** |  |
| **# of patients** | 40 | 24 | 147 | 618 |
| **Males/Females, *n*** | 26/14 | 17/7 | 81/66 | 319/299 |
| **Age (0-14), *n*** | 0 | 0 | 0 | 0 |
| **Age (15-39), *n*** | 2 | 3 | 27 | 196 |
| **Age (40-59), *n*** | 13 | 12 | 45 | 336 |
| **Age (>60), *n*** | 25 | 9 | 75 | 86 |
| **WBC count at presentation × 10<sup>3</sup>/μL (median, range)** | 10.55<br>(0.8-218.5) | 45<br>(1.5-153) | 20<br>(1-297) | N.A. |
| **Percentage of bone marrow blasts** | 63.7<br>(30-90) | 16.5<br>(0.4-57) | 72<br>(11-99) | 67<br>(6-98) |
| **Cytogenetic risk group, *n*** |  |  |  |  |
| ELN favorable | 3 (7.5%) | 6 (26.1%) | 17 (12%) | 204 (34%) |
| ELN intermediate | 5 (12.5%) | 7 (30.4%) | 96 (65%) | 284 (46%) |
| ELN adverse | 32 (80%) | 5 (21.7%) | 32 (22%) | 127 (19%) |
| N.A. | 0 | 5 (21.7%) | 2 (1%) | 3 (1%) |
| ***TP53* status** |  |  |  |  |
| Mutated | 40 | 2 | 13 | 14 |
| Wild type | 0 | 18 | - | - |
| Not tested / Not available | - | 4 | 134 | 604 |
| **Induction chemotherapy** |  |  |  |  |
| 7+3 | 5 | 2 | 113 | Cancer and Leukemia Group B treatment protocols 8525, 8923, 9621, 9720, 10201 and 19808 (25). |
| Fludarabine-based | - | 8 | - |  |
| Daunorubicin + Ara-C | 21 | 0 | - |  |
| MAV<sup>^^</sup> | 12 | 5 | - |  |
| HMA | - | 3 | 14 |  |
| Lenalidomide | 1 | - | 9 |  |
| Other | 1 | 6 | 11 |  |
| **Cohort-wide median OS (months from diagnosis)** | 5.06<br>(0.03-158.3) | 16.5<br>(0.3-57) | 15.5<br>(0.1-118.1) | 17.2<br>(0.03-224.1) |

<sup>^</sup>Cases of newly diagnosed non-promyelocytic AML with RNA-sequencing data and clinical annotation. OS = overall survival. <sup>^^</sup>Mitoxantrone, Ara-C and etoposide.

### Data sources for in silico analyses

The first data series, hereafter referred to as The Cancer Genome Atlas (TCGA) series, consisted of RNA-sequencing data (Illumina HiSeq2000) from 147 adult AML patients with complete cytogenetic, immunophenotypic and clinical annotation who were enrolled on Cancer and Leukemia Group B treatment protocols 8525, 8923, 9621, 9720, 10201 and 19808 (25). Thirteen patients had a documented *TP53* mutation. RNA and clinical data were retrieved from cBioPortal for Cancer Genomics (https://www.cbioportal.org/). Level 3 RSEM-normalized RNASeqV2 data was downloaded from TCGA and log_2_-transformed prior to analysis. No further pre-processing was applied. For mRNA expression data, cBioPortal for Cancer Genomics computes the relative expression of an individual gene and tumor specimen to the gene’s distribution in all samples that are diploid for the gene in question. The returned value (*z*-score) indicates the number of standard deviations away from the mean of expression in all other tumor samples. To ensure high stringency, a *z*-score threshold of ±2.0 was used in all analyses. The second data series (E-MTAB-3444), hereafter referred to as the HOVON series (26), was retrieved from Array Express and encompassed three independent cohorts of adults (≤60 years) with *de novo* AML. BM and blood samples were collected at diagnosis and were analyzed on the Affymetrix Human Genome U133 Plus 2.0 Microarray (26,27). Patients were treated with curative intent according to the Dutch-Belgian Hematology-Oncology Cooperative Group and the Swiss Group for Clinical Cancer Research (HOVON/SAKK) AML-04, -04A, -29, -32, -42, - 42A, -43 or -92 protocols (available at http://www.hovon.nl). Of the 618 patients, 14 had a documented *TP53* mutation.

### RNA isolation from bulk BM suspensions

RNA was isolated and processed as previously described (28). Approximately 100 ng per sample of RNA extracted from bulk BM aspirates were analyzed on the nCounter^®^ FLEX analysis system (NanoString Technologies, Seattle, WA) using the PanCancer IO 360™ mRNA panel (for research use only and not for use in diagnostic procedures). The reporter probe counts, i.e., the number of times the color-coded barcode for that gene is detected, were tabulated in a comma separated value format for data analysis with the nSolver™ software package (version 4.0.62) and nSolver Advanced Analysis module (version 2.0.115; NanoString Technologies). The captured transcript counts were normalized to the geometric mean of the housekeeping reference genes included in the assay and the code set’s internal positive controls. The relative abundance of immune cell types and immuno-oncology biological signatures were computed as previously published (29,30).

### Flow cytometry

Cells (0.5×10^6^) were aliquoted into 12×75mm tubes and were incubated with 5μL Human FcR Blocking Reagent (Miltenyi Biotec, Bergisch Gladbach, Germany), fluorochrome-conjugated monoclonal antibodies against PD-L1 (clone 29E.2A3) and HLA-A,B,C (clone W6/32; BioLegend, San Diego, CA, USA), and LIVE/DEAD fixable viability dyes (ThermoFisher Scientific, Waltham, MA, USA) for 30 minutes at 4°C, protected from light. Cells were finally re-suspended in 350 µL PBS and were run through a Gallios™ flow cytometer (Beckman Coulter, High Wycombe, UK). Data were analyzed with the Kaluza™ software package, v1.3 (Beckman Coulter).

### AML cell lines

For *in vitro* modeling experiments, commercial AML cell lines that harbor a missense (R248Q; Kasumi-1 cells; ATCC^®^ CRL-2724™) and a truncating mutation of *TP53* (KG-1 cells; ATCC^®^ CRL-246™), respectively, were selected. Kasumi-1 cells were cultured in RPMI (Lonza, Basel, Switzerland) supplemented with 20% fetal bovine serum (FBS; HyClone™; GE Healthcare Life Sciences, Pittsburgh, PA, USA) and 2 mM L-glutamine (Lonza). KG-1 cells were cultured with IMDM containing 25 mM HEPES and L-glutamine +20% FBS. Cells were seeded at 1.5×10^6^ per well in a 6-well plate with or without 100 IU IFN-γ (R&D systems, Bio-Techne Ltd., UK) and were harvested after 24 hours for further processing. Cell lysates of AML cell lines were used for the integrated measurement of mRNA, protein and single nucleotide variants (SNV) with the nCounter Vantage 3D™ Heme Panel (NanoString Technologies), as per manufacturer’s protocol. Cell lines HuT-78 (mature T-cells from a case of Sezary syndrome) and CCRF-CEM (T-cell acute lymphoblastic leukemia) with known mutations in key cancer drivers were used as controls.

### Gene ontology (GO) and gene set enrichment analysis (GSEA)

Metascape.org was used to enrich genes for GO biological processes and pathways. For the gene list submitted to metascape.org, pathway and process enrichment analyses are carried out using all genes in the genome as the enrichment background. Terms with a P value <0.01, a minimum count of 3, and an enrichment factor >1.5 (defined as the ratio between the observed counts and the counts expected by chance) are collected and grouped into clusters based on their membership similarities. GSEA was performed using the GSEA software v.3.0 (Broad Institute, Cambridge, USA) (31). Hallmark *TP53* oncogenic gene signatures (M2698 and M2694) were downloaded from the Molecular Signature Database (MSigDB). The analysis of functional protein association networks was performed using STRING (https://string-db.org/).

### Statistical analyses

Descriptive statistics included calculation of mean, median, SD, and proportions to summarize study outcomes. Comparisons were performed with the Mann-Whitney U test for paired or unpaired data (two-sided), as appropriate, or with the ANOVA with correction for multiple comparisons. A two-tailed *p* value <0.05 was considered to reflect statistically significant differences. The log-rank (Mantel-Cox) test was used to compare survival distributions. OS was computed from the date of diagnosis to the date of death. Relapse-free survival (RFS) was measured from the date of first CR to the date of relapse or death. Subjects lost to follow-up were censored at their date of last known contact. IBM SPSS Statistics (version 24) and GraphPad Prism (version 8) were used for statistical analyses.

## Results

### *TP53* mutational status correlates with immune infiltration in TCGA-AML cases

It has been shown that genetic drivers of solid tumors dictate neutrophil and T-cell recruitment, thus affecting the immune contexture and potentially assisting patient stratification (32). We first asked whether the expression of known AML drivers, including *TP53*, correlates with the immune composition and functional orientation of the BM tumor microenvironment (TME). To address this hypothesis, we retrieved RNA-sequencing data with cytogenetic and clinical annotation, including RFS and OS, from adult patients with non-promyelocytic AML (n=147 cases available through cBioPortal for Cancer Genomics; n=118 cases with information on prognostic molecular lesions). Patients had a median age of 60 years, 54% were male, with 12%, 65% and 22% classified as favorable, intermediate and adverse risk, respectively, based on 2017 European Leukemia-Net (ELN) risk stratification by genetics (**Table 1**). One hundred thirteen patients (77%) were reported as having received 7+3 cytotoxic induction chemotherapy. The remaining patients were treated with adjunctive therapy in addition to 7+3 or with hypomethylating agents (HMA). Immune signature scores were calculated as pre-defined linear combinations (weighted averages) of biologically relevant gene sets, as previously published (29,30). ELN intermediate cases with information on *NPM1* mutational status and *FLT3*-ITD were further subclassified into molecular low risk (*NPM1* mutations without *FLT3*-ITD) and molecular high risk cases (*NPM1* wild-type with *FLT3*-ITD) (33). *TP53* mutations (11 missense, 4 frameshift and 4 splice site) were present in 13 patients (**Supplemental Table 1**).

As shown in **Fig. 1A**, p53 mutated AML cases showed higher levels of immune infiltration compared with patients with low-risk or intermediate-risk molecular lesions and with patients harboring other high-risk molecular features (*RUNX1* mutations and *NPM1* wild-type with FLT3-*ITD*). p53 mutated cases had higher TMB relative to TCGA-AML cases without any known *TP53* abnormality (**Fig. 1B**). The tumor inflammation signature (TIS) score, an established predictor of response to immune checkpoint blockade (ICB) across a broad range of solid tumors (34,35), was significantly higher in *TP53* mutated cases relative to cases with molecular lesions associated with favorable clinical outcomes (*NPM1* mutations without *FLT3*-ITD) and to cases with clonal hematopoiesis of indeterminate potential (CHIP)-defining mutations (*AXSL1, TET2*, and *DNMT3A*; **Fig. 1C**). Other gene expression scores reflecting an immune-infiltrated TME (23), such as the IFN-γ signaling, inflammatory chemokine and lymphoid scores, were significantly higher in *TP53* mutated cases (**Fig. 1C**), suggesting a higher degree of lymphoid infiltration and the activation of IFN-γ-related signaling pathways. Interestingly, *TP53* mutated cases had higher expression of immune checkpoints (*PD-L1* and *TIGIT*) and molecules reflecting a highly immunosuppressive TME, such as the Treg-associated transcription factor *FOXP3* (**Fig. 1D**). Survival estimates for TCGA-AML cases with mutated *TP53* are summarized in **Fig. 1E** which shows that median OS from diagnosis was 4.5 months compared with 16.3 months in patients with other prognostic molecular lesions (detailed in **Fig. 1A**).

**Figure 1:**
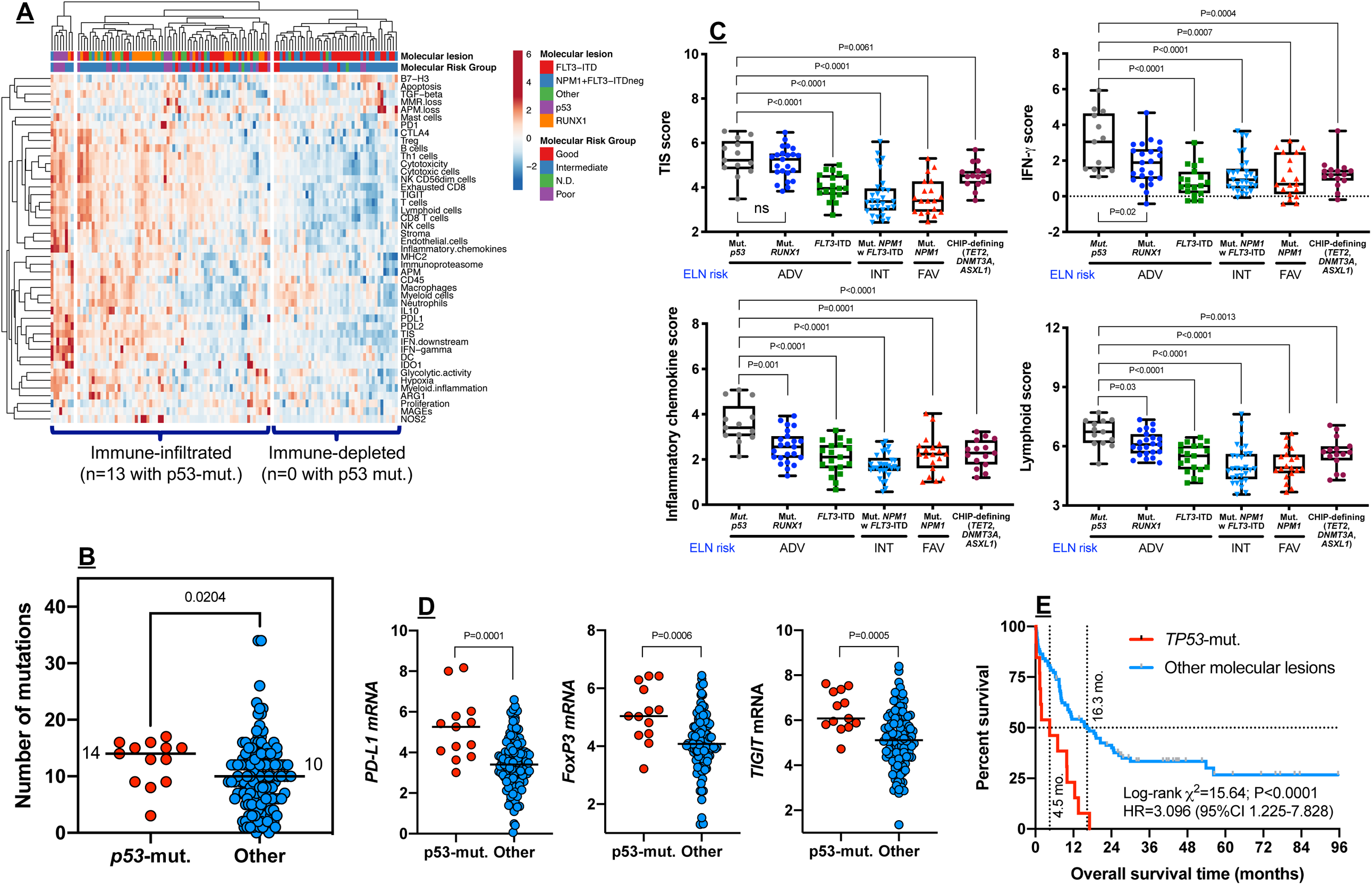
*TP53* mutations correlate with an immune-infiltrated TME in TCGA-AML. **A**) Heat-map of immune cell type-specific scores and biological activity scores in TCGA-AML cases with information on prognostic molecular lesions (n=118; unsupervised hierarchical clustering; Euclidean distance; complete linkage). ClustVis, an online tool for clustering of multivariate data, was used for data analysis and visualization (63). *NPM1* = nucleophosmin-1; *FLT3*-ITD = fms-like tyrosine kinase 3 internal tandem duplication. European Leukemia-Net (ELN) intermediate cases were further subclassified into molecular low risk (*NPM1* mutations without *FLT3*-ITD) and molecular high risk cases (*NPM1* wild-type with *FLT3*-ITD) (33). **B**) Tumor mutational burden (TMB) in TCGA-AML cases with *TP53* mutations or with other prognostic molecular lesions. Bars denote median values. Data were compared using the Mann-Whitney *U* test for unpaired determinations. **C**) Box plots showing immune signature scores in TCGA-AML cases with *TP53* mutations and other prognostic molecular lesions. TIS = tumor inflammation signature. Data were compared using the Kruskal-Wallis test for unpaired determinations. **D**) Expression of *FoxP3* and negative immune checkpoints *PD-L1* and *TIGIT* in TCGA-AML cases with *TP53* mutations and other prognostic molecular lesions. Bars denote median values. Data were compared using the Mann-Whitney *U* test for unpaired determinations. **E**) Kaplan-Meier estimate of overall survival in TCGA-AML cases with *TP53* mutations and with other prognostic molecular lesions, as defined above. Survival curves were compared using a log-rank test. HR = hazard ratio.

### Primary BM samples from patients with *TP53* mutated AML express inflammatory and IFN-related gene sets

We next compared immune gene expression profiles between bulk BM specimens from patients with *TP53* mutated AML (n=42) and *TP53* wild type AML (n=22). The predicted functional consequences of p53 mutations (84% missense; **Supplemental Fig. 1A**) are listed in **Supplemental Table 2** (36). Pre-defined immune cell type-specific scores and biological activity scores (30) distinguished patients with *TP53* mutated AML from individuals with *TP53* wild type AML, as highlighted by principal component analysis (**Fig. 2A**). The frequency of *TP53* mutated cases was 87% (20/23), 75% (21/28) and 8% (1/13) in patients with high, intermediate and low levels of immune infiltration, respectively (**Fig. 2B**). We next analyzed the immune transcriptomic profile at the gene level and identified a set of 34 differentially expressed (DE) immune genes at a false discovery rate (FDR) <0.01 between patients with *TP53* mutated AML and *TP53* wild type AML (**Fig. 2C-D** and **Supplemental Table 3**) which will be herein referred to as *TP53* immune gene classifier. The *TP53* immune signature genes have not been previously implicated in the *TP53* pathway, as shown in **Supplemental Fig. 1B**. Neutrophil chemoattractants (pro-inflammatory *CXCL1, CXCL2* and *CXCL8* or *IL8*) and IFN-inducible molecules such as *CCL2, IL33, IL6, OASL* and *RIPK2* were more highly expressed in *TP53* mutated compared with *TP53* wild type patients (**Fig. 2D**). The pattern recognition scavenger receptor *MARCO*, which defines tumor-associated macrophages with an M2-like immunosuppressive signature in experimental tumor models (37) and in patients with lung cancer (38), was more abundant in *TP53* mutated AML. Furthermore, *TP53* mutated AML expressed significantly higher levels of *TP53* pathway genes p21 (*CDKN1A*) and *Fas* (**Fig. 3A**), as well as *IFNG, FOXP3, PD-L1, LAG3, CD8A* and *GZMB*, a molecule recently associated with features of exhaustion and senescence in AML infiltrating CD8^+^ T cells (**Fig. 3B**) (39). The DE molecules exhibited enrichment of gene ontologies (GO) and KEGG pathways related to inflammatory responses, cellular response to cytokine stimuli, response to stress, cytokine-cytokine receptor interactions and IL-17-mediated and TNF-mediated signaling (**Fig. 3C** and **Supplemental Table 4**). We next computed scores that capture frequently dysregulated signaling pathways in cancer using pre-defined sets of relevant genes. As shown in **Supplemental Fig. 1C**, *TP53* mutated cases expressed higher levels of NF-κB, JAK/STAT and PI3K-Akt signaling molecules relative to BM samples from patients with *TP53* wild-type AML. In contrast, DNA damage repair genes as well as Hedgehog and Wnt signaling pathway genes were upregulated in *TP53* wild type AML compared with *TP53* mutated AML (**Supplemental Fig. 1C-D**). These findings are congruent with previous studies showing that *TP53* is a suppressor of canonical Wnt signaling in solid tumors (40).

**Table 2:**
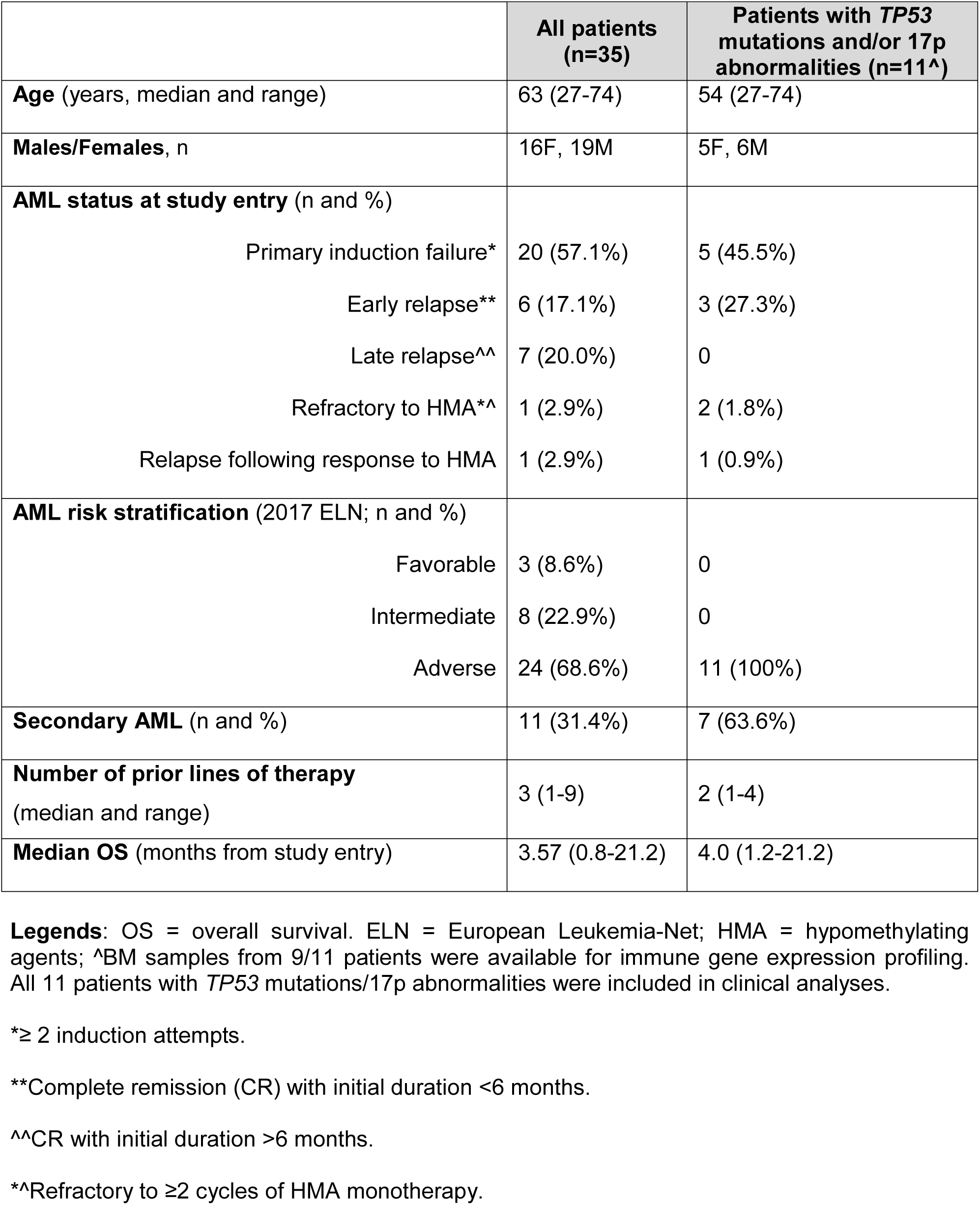
Patients’ characteristics (immunotherapy cohort)

**Figure 2:**
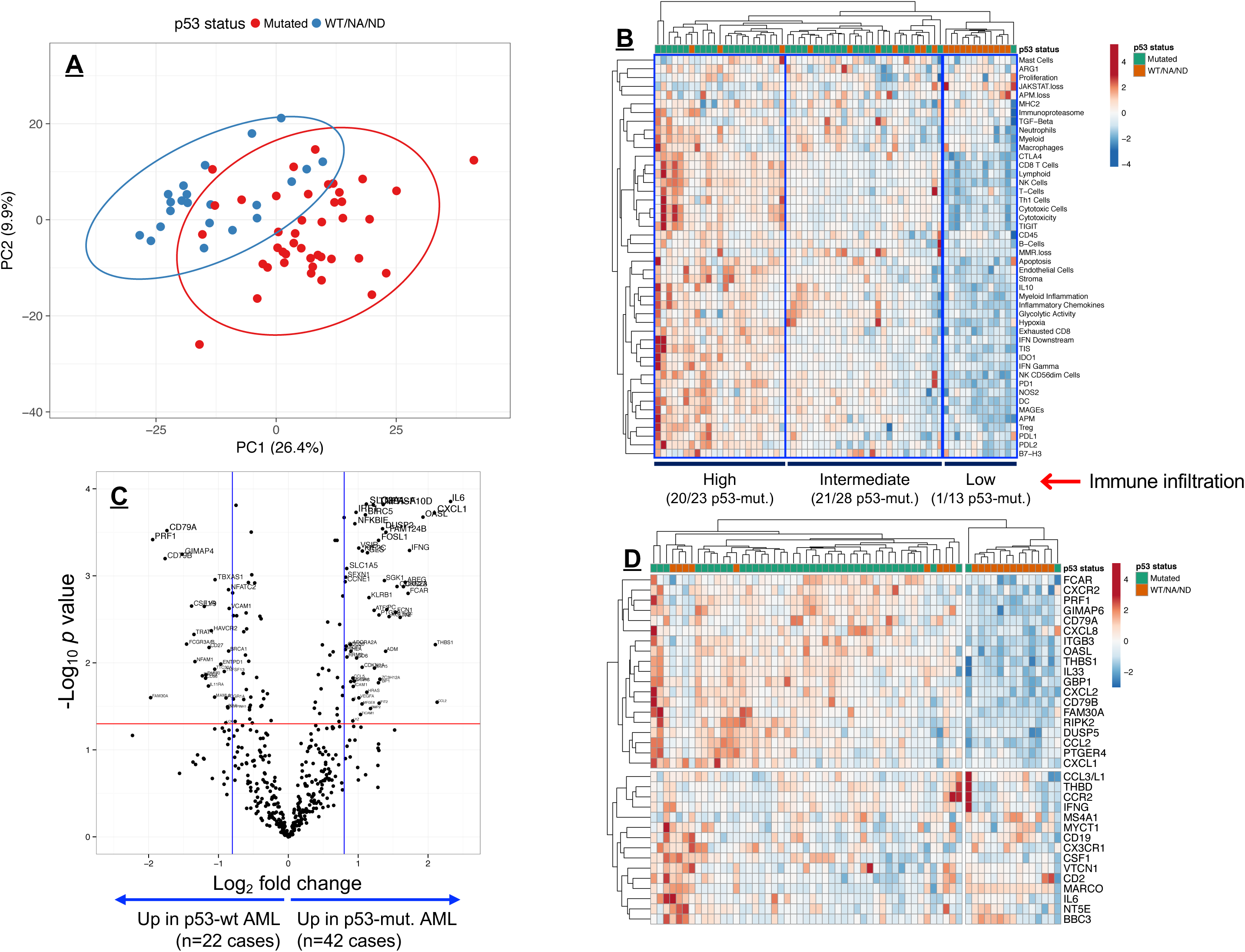
Identification of a *TP53* immune gene classifier in patients with *TP53*-mutated AML (SAL and Bologna cohorts). **A**) Principal component analysis (PCA) of 770 immune genes (IO 360 Panel) in patients with *TP53* mutated AML (n=42) and *TP53* wild type AML (n=22). Points are colored by p53 mutational status (mutated = red; wild type = blue). ClustVis was used for data analysis and visualization. **B**) Heatmap of immune cell type-specific and biological activity scores in patients with *TP53* mutated AML and *TP53* wild type AML (unsupervised hierarchical clustering; Euclidean distance; complete linkage). The number of *TP53* mutated cases in each immune cluster (high, intermediate, low) is indicated. ClustVis, an online tool for clustering of multivariate data, was used for data analysis and visualization. ND = Not determined; NA = Not available. **C**) Volcano plot showing differentially expressed genes between patients with *TP53* mutated AML and *TP53* wild type AML. Plots were drawn using an online server hosted on shinyapps.io by RStudio (https://paolo.shinyapps.io/ShinyVolcanoPlot/). **D**) Heatmap of differentially expressed (DE) genes between patients with *TP53* mutated AML and *TP53* wild type AML (P value threshold = 0.01; log_2_ fold change ≥1.5-fold). ClustVis, an online tool for clustering of multivariate data, was used for data analysis and visualization.

**Figure 3:**
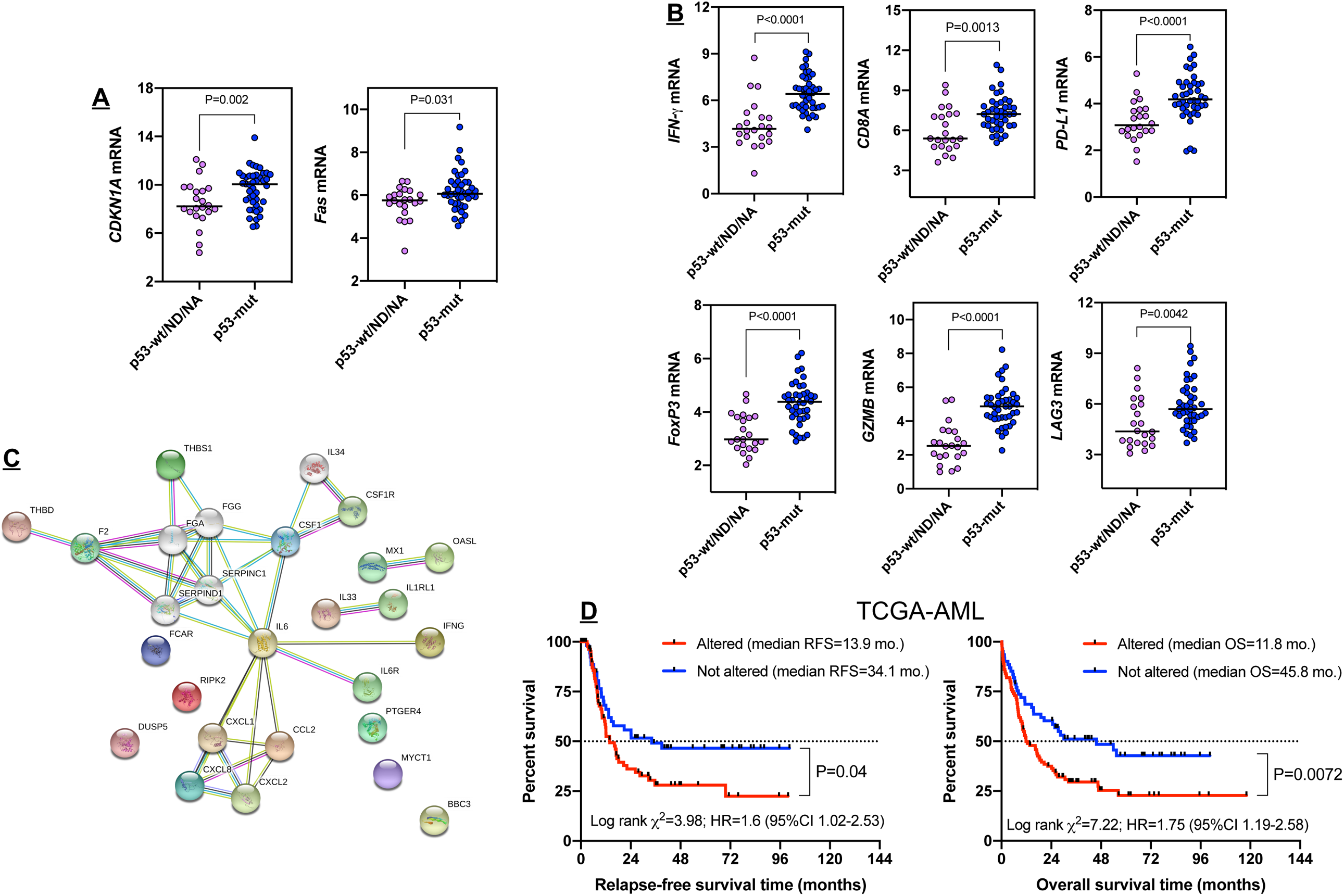
Identification of a *TP53* immune gene classifier in patients with *TP53*-mutated AML (SAL and Bologna cohorts). **A**) Expression of *TP53*-inducible genes p21 (*CDKN1A*) and *Fas* in patients with *TP53* mutated and *TP53* wild type AML. Data were compared using the Mann-Whitney *U* test for unpaired determinations. **B**) Box plots summarizing the expression levels of negative immune checkpoint and immune genes related to T-cell infiltration, regulatory T cells and cytolytic activity in patients with *TP53* mutated and *TP53* wild-type AML. Bars denote median values. Data were compared using the Mann-Whitney *U* test for unpaired determinations. **C**) Analysis of functional protein association networks using STRING (https://string-db.org/). Top 10 molecules interacting with DE genes in the *TP53* immune classifier are shown together with their predicted mode of action (highest confidence interaction scores >0.900). Network nodes (query proteins) represent proteins produced by a single protein-coding gene locus. White nodes represent second shells of interactors. Empty and filled nodes indicate proteins of unknown or partially known 3-dimensional structure, respectively. Edges represent protein–protein associations. Line shapes denote predicted modes of action. **D**) Abnormalities in *TP53* immune classifier genes (mRNA upregulation, gene amplification, deep deletion and mis-sense mutations, relative to the gene’s expression distribution in all profiled AML samples) were detected in 101 of 162 (62%) sequenced BM samples from TCGA. Data were retrieved, analyzed and visualized using cBioPortal. Abnormalities in only one gene utilized in the cBioPortal query (by default, non-synonymous mutations, fusions, amplifications and deep deletions) were sufficient to define that particular patient sample as “altered”. The Kaplan-Meier method was used to generate survival curves, which were compared using a log-rank test. RFS = relapse-free survival; OS = overall survival; HR = hazard ratio.

The *TP53* immune gene classifier that we identified in primary AML BM samples was further assessed *in silico* for potential prognostic value in TCGA-AML cases. Abnormalities of the 34 DE immune genes (including mRNA up-regulation, amplification, deep deletion and mis-sense mutations) significantly correlated with *TP53* mutational status (P=9.95×10^−3^), with higher levels of immune infiltration, and with the expression of negative immune checkpoints and IFN signaling molecules (**Supplemental Fig. 2A-C**). Importantly, RFS and OS estimates were significantly worse for TCGA-AML patients with abnormalities in query genes (**Fig. 3D**). Taken together, these findings suggest that the immunological TME of *TP53* mutated AML is inherently pro-inflammatory and IFN-γ-dominant, and that these molecular features correlate with poor clinical outcomes.

### Loss-of-function (LOF) *TP53* mutations correlate with enhanced IFN-γ and inflammatory signaling in AML cell lines

It has been reported that LOF is frequent among *TP53* missense mutations (41). The consequences of mutant p53 expression on IFN signaling have not been evaluated previously. We therefore performed *in vitro* modelling experiments with commercial AML cell lines with known p53 GOF/LOF status. DNA single nucleotide variant (SNV) amplicons, mRNA and protein lysates were prepared as detailed in Materials and Methods. The inter-assay reproducibility of mRNA and protein measurements is shown in **Supplemental Fig. 3A**. The SNV assay confirmed the presence of *FBXW7* (R465C), *KRAS* (G12D) and *MLH1* (I219V) and *TP53* (R248Q) mutations in HuT-78 cells (data not shown), in accordance with available knowledge from the COSMIC database (https://cancer.sanger.ac.uk/cosmic). Similarly, we detected known mutations in *JAK3* (A573V), *MLH1* (I219V), *NRAS* (Q61K) and *TP53* (R196*) in control CCRF-CEM cells (data not shown), providing an *in-silico* validation of the NanoString SNV assay. KG-1 cells harbored a sequence change (c.672+1G>A) that affects a donor splice site in intron 6 of the *TP53* gene, resulting in a loss of protein function. As expected, TP53 protein was undetectable in KG-1 cells, with less than 10 log_2_ fold-change compared with the Kasumi-1 AML cell line; **Supplemental Fig. 3B**). A known GOF mutation in *TP53* (R248Q) (42) was detected in Kasumi-1 AML cells (**Supplemental Fig. 3B**).

A list of genes was generated by considering the FDR (<0.01) and fold change (ranging from -1.7 to 1.7) of genes that were differentially expressed between KG-1 and Kasumi-1 cells (**Fig. 4A**). Specifically, KG-1 AML with a LOF *TP53* mutation over-expressed genes involved in IFN-mediated signaling and inflammation, including *HGF, CIITA, PIM1, OSM, STAT1* and *IRF1* (**Fig. 4B** and **Supplemental Table 5**). Furthermore, KG-1 cells showed higher expression of PD-L1 and class I molecules, which are known to be regulated by IFN-γ, compared with Kasumi-1 cells (**Supplemental Fig. 3C**). PI3K-Akt, NF-κB, JAK/STAT and *TP53* pathway genes, as well as genes associated with T helper 17 (Th17) differentiation, were significantly enriched in KG-1 cells, as shown in **Fig. 4C** and in line with our findings in patients with *TP53* mutated AML (**Supplemental Fig. 1B-C**). GO and KEGG pathways captured by the DE genes between KG-1 and Kasumi-1 AML are listed in **Supplemental Table 6**. Finally, **Fig. 4D** summarizes the analysis of functional protein association networks and shows the top 10 molecules interacting with DE genes.

**Figure 4:**
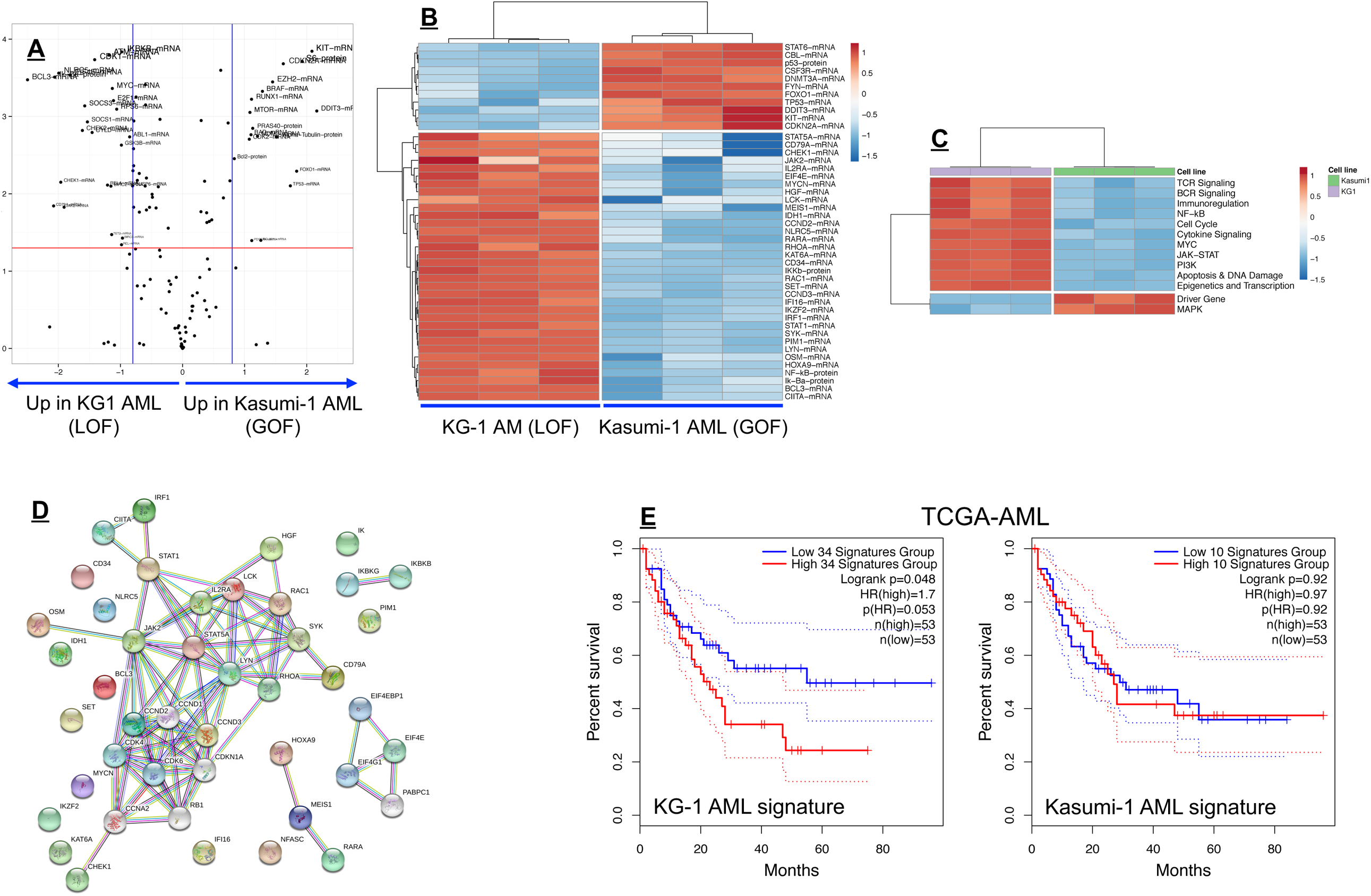
Integrated mRNA and protein profile in AML cells lines with gain-of-function (GOF) and loss-of-function (LOF) *TP53* mutations. **A**) Volcano plot showing differentially expressed (DE) mRNA species and proteins between AML cell lines with GOF (Kasumi-1; p.R248Q; Broad Institute Cancer Cell Line Encyclopedia) (64) and LOF (splice site) mutations of *TP53*. Plots were drawn using an online server (https://paolo.shinyapps.io/ShinyVolcanoPlot/) hosted on shinyapps.io by RStudio. **B**) Heat-map of the top DE mRNA species and proteins between KG-1 AML and Kasumi-1 AML (unsupervised hierarchical clustering; Euclidean distance; complete linkage). ClustVis, an online tool for clustering of multivariate data, was used for data analysis and visualization (63). **C**) Heat-map of signaling pathway scores in KG-1 AML and Kasumi-1 AML (unsupervised hierarchical clustering; Euclidean distance; complete linkage). **D**) Analysis of functional protein association networks using STRING (https://string-db.org/). Top 10 molecules interacting with DE mRNAs and proteins between KG-1 AML and Kasumi-1 AML are shown together with their predicted mode of action (highest confidence interaction scores >0.900). Network nodes (query proteins) represent proteins produced by a single protein-coding gene locus. White nodes represent second shells of interactors. Empty and filled nodes indicate proteins of unknown or partially known 3-dimensional structure, respectively. Edges represent protein–protein associations. Line shapes denote predicted modes of action. **E**) Kaplan-Meier (KM) estimate of survival from diagnosis in TCGA-AML cases with abnormalities in DE expressed genes between KG-1 (n=34) and Kasumi-1 AML cells (n=10). KM curves (median split of signature scores) were generated using GEPIA2, an enhanced web server for TCGA gene expression profiling and interactive analysis (http://gepia2.cancer-pku.cn/#index) (65). Signature scores are calculated as the mean value of log_2_ transcripts per million (TPM). GEPIA2 uses the log-rank (Mantel-Cox) test to compare survival distributions. HR = hazard ratio.

We next assessed whether the experimentally derived gene/protein signatures could be of potential significance for survival prediction in TCGA-AML cases. As shown in **Fig. 4E**, genes/proteins overexpressed in KG-1 AML (n=34; **Supplemental Table 5**) predicted for significantly shorter OS (log-rank P value=0.048). In contrast, genes upregulated in Kasumi-1 AML were unable to stratify patient survival (**Fig. 4E**). Overall, these experiments suggest that LOF *TP53* mutations correlate with heightened IFN-γ signaling and activation of other intracellular signaling pathways, including PI3K-Akt, JAK/STAT and NF-κB, and that the above molecular features may correlate with worse clinical outcomes.

### *TP53* mutated patients with relapsed/refractory AML show evidence of anti-leukemic activity of flotetuzumab immunotherapy

We have previously shown that baseline IFN-γ-related mRNA profiles, including the TIS score, are associated with response to flotetuzumab, a CD3×CD123 DART^®^ molecule, in R/R AML (43,44). The heightened expression of IFN-γ pathway molecules that we observed in *TP53* mutated AML suggests that this patient subset may also benefit from T-cell engaging immunotherapies, such as flotetuzumab. To test this hypothesis, we correlated *TP53* mutational status with immune landscapes and with anti-leukemic activity (ALA) from flotetuzumab in a cohort of 35 patients with R/R AML treated with flotetuzumab. Patients’ characteristics, including *TP53* mutational status and/or the presence of chromosome 17p deletions usually associated with loss of one allele of *TP53* and mutation/loss of the other (45), are summarized in **Table 2**. Baseline BM samples for immune gene expression profiling were available in 9/11 patients with *TP53* mutations and/or genomic loss of *TP53*; among these, 7/9 patients showed high or intermediate levels of immune infiltration (**Fig. 5A**). Overall, ALA, which was defined as >30% reduction of BM blasts from baseline, was documented in 45.5% (5 out of 11) evaluable patients with *TP53* mutations and/or 17p abnormalities (2 CR, 1 CRh, 1 MLFS, and 1 OB). Time on treatment and time to patient death and/or censoring are summarized in **Fig. 5B** for individuals with *TP53* mutations and/or 17p deletion, including two patients who proceeded to receive allogeneic HSCT. The reduction of BM blasts in 10 patients with *TP53* abnormalities for whom a post-cycle 1 BM sample was available averaged 42% (**Fig. 5C**). In p53 mutated patients with evidence of ALA, the TIS, inflammatory chemokine, Treg and IFN-γ gene expression scores were significantly higher at baseline compared with non-responders (**Fig. 5D**), highlighting the association between response to T-cell engagers and a T cell inflamed and highly immunosuppressed TME (43). Median OS from study entry was 4.0 months (range 1.25-21.25) for patients with *TP53* abnormalities (**Fig. 5E**), indicating that flotetuzumab immunotherapy may alleviate the negative prognostic impact of *TP53* mutations. The survival estimates for patients with *TP53* mutated AML treated with flotetuzumab compare favorably with survival predictions for *TP53* mutated cases with PIF (median OS=1.16 months) in large AML series, such as the HOVON cohort (**Fig. 6A**). *In silico* analyses also suggest that median OS is not dissimilar between newly diagnosed HOVON cases with *TP53* mutations (13 patients; 3.58 months) and with PIF (125 patients; 3.78 months; **Fig. 6B**). Finally, gene set enrichment analysis (GSEA) with all transcripts in the HOVON dataset provided as input and ranked by the log_2_ fold-change between chemotherapy non-responders (PIF) and responders showed the increased expression of a curated hallmark set of 172 genes linked to the *TP53* pathway in patients with PIF (**Fig. 6C**).

**Figure 5:**
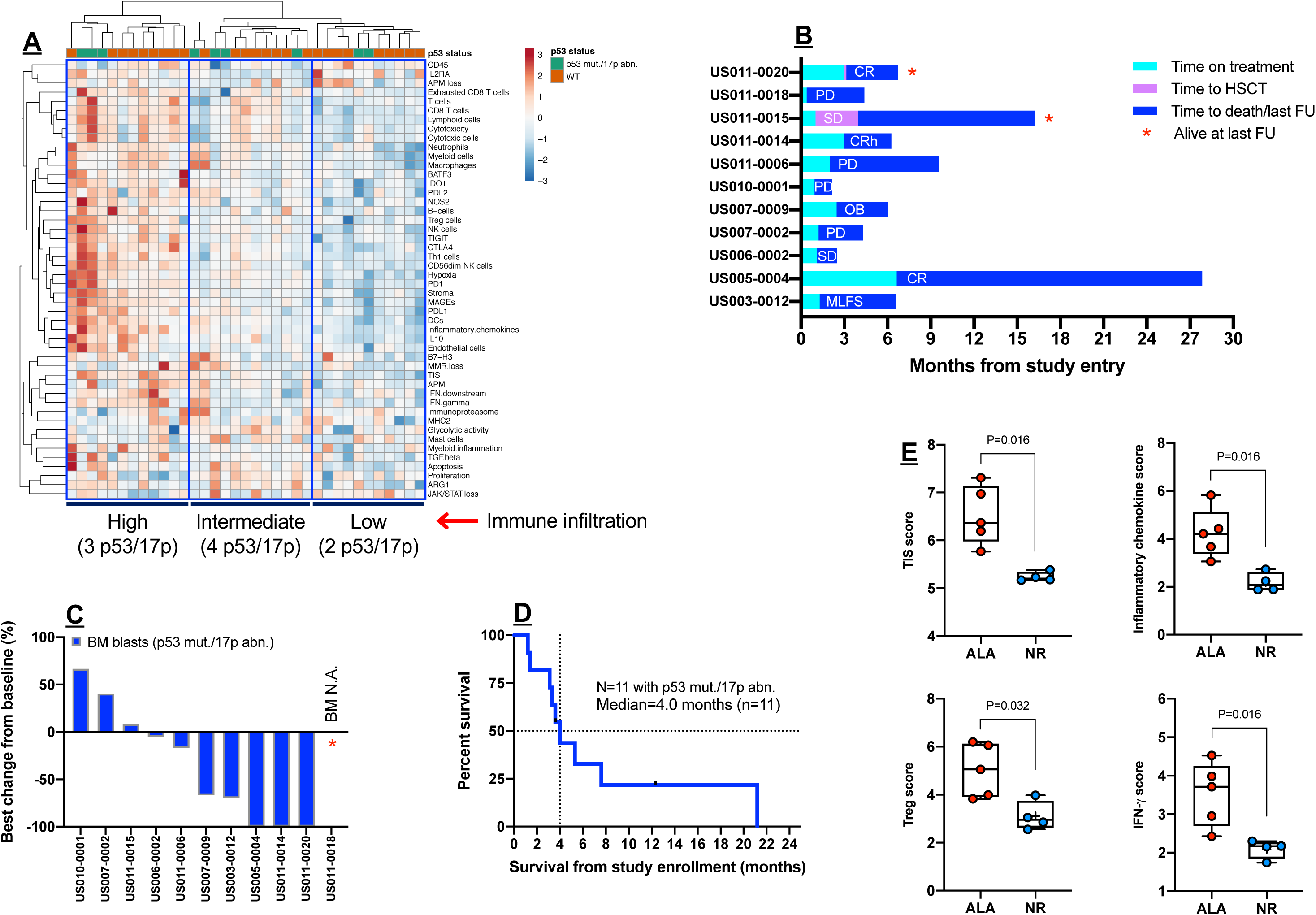
Immune landscape and immunotherapy response in patients with relapsed/refractory AML with *TP53* mutation and/or 17p abnormalities. **A**) Heat-map of immune cell type-specific scores and biological activity scores in AML patients treated with flotetuzumab immunotherapy. ClustVis, an online tool for clustering of multivariate data, was used for data analysis and visualization (63). **B**) Time on treatment and time to death and/or censoring in relation to clinical responses in the 11 patients with *TP53* mutations and/or 17p deletions. CR = complete remission; CRh = complete remission with partial hematopoietic recovery; SD = stable disease; OB = other benefit; PD = progressive disease; MLFS = morphological leukemia-free state. Response criteria were described in materials and Methods. FU = follow-up. **C**) Waterfall plot depicting changes in BM blasts after cycle 1 of flotetuzumab in patients with *TP53* mutation and/or 17p deletion with genomic loss of *TP53* (n=10). A BM sample was not available in 1 patient who progressed on treatment. **D**) Kaplan-Meier estimate of survival from study entry in patients with *TP53* mutations and/or 17p deletions receiving flotetuzumab immunotherapy. **E**) Tumor inflammation signature (TIS) score, inflammatory chemokine score, regulatory T-cell (Treg) score and IFN-γ score in baseline bone marrow (BM) samples from patients with *TP53* mutations and/or 17p deletion with genomic loss of *TP53*. ALA = anti-leukemic activity; NR = non-responder. Data were compared using the Mann-Whitney *U* test for unpaired determinations.

**Figure 6:**
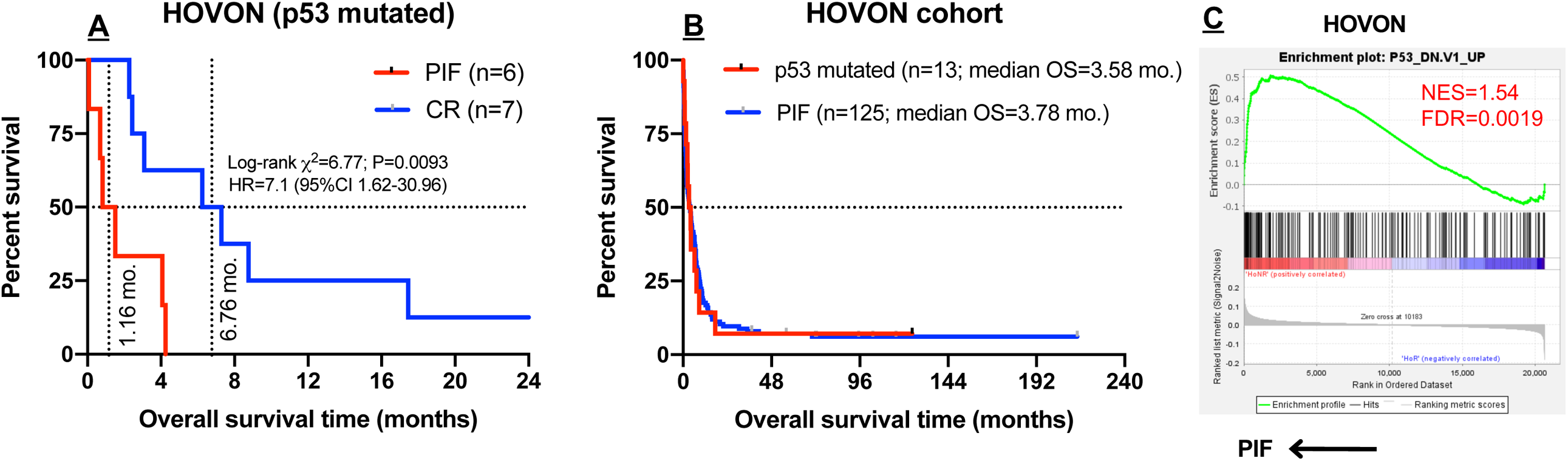
*TP53* pathway genes are enriched in HOVON AML cases with primary induction failure. **A**) Overall survival time in patients with *TP53* mutated AML (HOVON cohort) with primary induction failure (PIF; n=6) and with complete response (CR; n=6) to induction chemotherapy. The log-rank (Mantel-Cox) test was used to compare survival distributions. HR = hazard ratio. **B**) Kaplan-Meier estimate of overall survival (OS) in HOVON cases with *TP53* mutations (n=13) and PIF (n=125). The log-rank (Mantel-Cox) test was used to compare survival distributions. **C**) Gene set enrichment analysis (GSEA) with all transcripts in the HOVON dataset provided as input and ranked by the log_2_ fold-change between chemotherapy non-responders (primary induction failure, PIF) and responders. GSEA was performed using the GSEA software v.3.0 (Broad Institute, Cambridge, USA) (31). A *TP53* pathway gene set [n=172 genes upregulated in National Cancer Institute (NCI)-60 cancer cell lines with mutated p53 (M2698)] was downloaded from the Molecular Signature Database (MSigDB) (29,30). NES = normalized enrichment score; FDR = false discovery rate (adjusted p value); PIF = primary induction failure.

## Discussion

This multi-cohort study provides evidence for a correlation between IFN-γ-dominant immune subtypes of AML and *TP53* abnormalities by showing that *TP53* mutated cases exhibit higher levels of CD8^+^ T-cell infiltration and IFN-γ signaling compared with AML subgroups with other risk-defining molecular lesions, including *RUNX1, ASXL1* and CHIP-related mutations. Previously described gene expression-based predictors of response to ICB in solid tumors, such as the TIS (34,35), as well as negative immune checkpoints *PD-L1, TIGIT* and *LAG3* were significantly more expressed in AML with *TP53* mutations relative to other molecular subtypes. Some of the characterized genes may therefore contribute mechanistically to the poor prognosis associated with *TP53* mutated AML through the induction of immune escape. By comparing immune gene expression profiles between primary BM samples from patients with *TP53* mutated and *TP53* wild type AML, we identified a 34-gene immune classifier that is enriched in gene ontologies related to IFN-γ/inflammatory responses and IL-17/TNF-mediated signaling and that stratifies RFS and OS in a large cohort of TCGA-AML cases. Similar to previous studies that computed an expression signature of *TP53* mutated breast cancer (46), our *TP53* immune classifier genes showed no overlap with known *TP53* pathway genes.

Recent evidence supports a novel role for p53 in regulating immune responses and inflammation, in addition to its well-characterized function as a tumor suppressor (15). In mice infected with influenza virus, *TP53* directly activates expression of immune response genes, including IFN-inducible molecules such as *IRF5, IRF9* and *ISG15* (47). Mice harboring a germline *TP53* mutation develop severe chronic inflammation with failure to resolve tissue damage, and are highly susceptible to develop inflammation-associated colon cancer, suggesting *in vivo* pro-inflammatory and immune-related GOF (48). Human cancer cells with a GOF mutation of *TP53* can reprogram macrophages to a tumor-supportive and anti-inflammatory phenotype with increased activity of TGF-β (49). Intriguingly, colorectal cancer patients with GOF mutations of *TP53* (i.e., positions R245, R248, R175, R273, R282) have dense tissue infiltration with CD206-positive tumor-associated macrophages, over-express inflammatory and oncogenic gene signatures, and experience shorter OS (49). A recent analysis has suggested a correlation between loss of *TP53* function and absence of a cytotoxic T lymphocyte gene signature in estrogen receptor-negative breast cancer, leading to failure of tumor immunosurveillance (50). Recent *in silico* analyses of METABRIC and TCGA breast cancers have shown that *TP53* mutated tumors display higher expression of lymphocytic and cytotoxicity markers, *STAT1*, molecules implicated in antigen processing and presentation, as well as activation of JAK/STAT signaling (51). Counterintuitively, higher expression of negative immune checkpoints, Treg cell signatures and metastasis-promoting genes correlated with longer OS times in *TP53* mutated patients (51).

*In vitro* modeling experiments using commercial AML cell lines with LOF/GOF mutations of *TP53* allowed us to identify a set of DE mRNAs/proteins between KG-1 cells (*TP53* truncating mutation) and Kasumi-1 cells (p.R248Q GOF mutation), and suggested that *TP53* LOF, which is frequent among *TP53* missense mutants (52), may translate into the upregulation of IFN pathway molecules, Th17 genes, and intermediates involved in JAK/STAT and PI3K-Akt signaling and in the pro-inflammatory NF-κB pathway. Notably, abnormalities of the DE mRNA/proteins in the KG-1 AML signature correlated with higher BM immune infiltration and with *TP53* mutations in TCGA cases and were prognostic, as suggested by the significant separation of the survival curves. In sharp contrast, genes/proteins overexpressed in Kasumi-1 cells harboring a GOF mutation of *TP53* were unable to stratify survival in TCGA cases.

Pharmacological *TP53* reactivation is actively being pursued in patients with AML. MDM2 is an E3 ubiquitin ligase that binds to *TP53* and induces its proteasomal degradation. Treatment with DS-5272, an inhibitor of the *TP53*-MDM2 interaction, in mouse models of AML was associated with the up-regulation of inflammatory and IFN-associated genes, including *PD-L1*, and translated into enhanced anti-leukemia control (53). Furthermore, the survival benefit provided by *TP53* reactivation with DS-5272 was largely mediated by NK cells (53). In a mouse model of tumor senescence, *TP53* restoration caused liver tumor cells to secrete NK cell-recruiting chemokines, including CCL2, CXCL1 and CCL5, therefore favoring tumor rejection (54). Whether *TP53* reactivation in patients with cancer is associated with immune-mediated therapeutic effects remains to be established in future clinical trials. Interestingly, 40% of patients with solid tumors display T-cell responses to *TP53* hotspot mutations which are mediated by both CD4^+^ and CD8^+^ T cells (55), underpinning the broad immunogenicity of *TP53* neoepitopes. In this respect, antigen-experienced T cells have been expanded *ex vivo* from nine patients with metastatic epithelial cancers expressing a hotspot *TP53* mutation and have been successfully screened for neoantigen responses (56). This observation raises the hypothesis that the presence of immunogenic *TP53* mutations accounted for the higher degree of immune infiltration and activation in our patient cohorts with *TP53* mutated AML, as also suggested by studies in solid tumors (20,57). Furthermore, the results presented here point to the establishment of an inherently immuno-suppressive and IFN-γ-driven TME in patients with *TP53* mutated AML, who might require combinatorial immunotherapy approaches that also target Treg cells and negative immune checkpoints, either concomitantly or sequentially.

Analyses of clinical outcomes in large public cohorts of patients with AML show OS estimates for patients with *TP53* mutations/17p abnormalities ranging from 3.58 months (HOVON series) to 4.5 months (TGCA series). These survival predictions are significantly worse than the 16.3 month OS estimate for patients with other molecular abnormalities (TCGA series), but similar to that of PIF AML patients without *TP53* abnormalities (3.78 months, HOVON series). AML PIF patients with *TP53* altered status, however, survived a median of only 1.16 months, further underlining the profound negative prognostic role of *TP53* abnormalities in AML.

Post hoc analyses of a cohort of 35 patients with relapsed/refractory AML treated with flotetuzumab suggested that immunotherapy may be efficacious in individuals with altered *TP53* status, with an overall reduction of BM blasts averaging 42% and with evidence of ALA in 45.5% (5/11) of the patients. Responders showed intermediate-to-high levels of BM immune infiltration at baseline and higher TIS, inflammatory chemokine and CD8 T-cell scores compared with non-responders, suggesting that the presence of *TP53* abnormalities does not hamper the response to immunotherapy providing an inflammatory gene signature is present. Preliminary survival data in R/R patients with *TP53* abnormalities that responded to flotetuzumab indicate a median OS of 4.5 months, further suggesting that immunotherapy may be beneficial in AML with *TP53* abnormalities, even in individuals with PIF. Interestingly, HOVON cases with PIF showed significant enrichment of *TP53* pathway genes, irrespective of their *TP53* mutational status, and had similar survival predictions to those in *TP53* mutated cases. These results help to shed light into the heterogeneity of molecular mechanisms underpinning resistance to intensive induction chemotherapy (58) and are congruent with previous studies showing that non-mutational and non-deletional *TP53* inactivation, leading to protein stabilization, overexpression of *MDM2* (a canonical negative regulator of *TP53*) and absence of p21 (a *TP53* pathway effector molecule) are highly prevalent in AML (59).

Other molecularly defined subtypes of AML, including AML with recurrent somatic mutations in the epigenetic regulator DNA methyltransferase 3A (*DNTM3A*) (60) and AML with *FLT3*-ITD, might benefit from immune interventions. It has been shown that Dnmt3a-mediated *de novo* methylation programming in tumor-infiltrating, PD1^high^ CD8^+^ T cells promotes functional T-cell exhaustion and represents a barrier to T-cell rejuvenation, thereby restricting the efficacy of experimental ICB therapy (61). Sorafenib treatment might favor metabolic reprogramming of human CD8^+^ T cells, inducing molecular features of longevity through the activation of the IRF7-IL-15 axis in leukemia cells, leading to eradication of *FLT3*-ITD^+^ AML (62).

In conclusion, our study shows that *TP53* mutations are associated with higher T-cell infiltration, expression of negative immune checkpoints and IFN-γ-driven transcriptional programs, and correlate with disease control in response to flotetuzumab immunotherapy. The anti-leukemic activity with flotetuzumab validates the translational relevance of our findings and encourages further studies of T-cell targeting immunotherapeutic approaches in patients with *TP53* mutations.

## Author contributions

### Concept and design

J.K. Davidson-Moncada, S. Rutella

### Development of methodology

J. Vadakekolathu, S. Reeder, S.E. Church, T. Hood, S. Rutella

### Acquired, consented and managed patients; processed patient samples

I. Aldoss, J. Godwin, M.J. Wieduwilt, M. Arellano, J. Muth, F. Ravandi, K. Sweet, H. Altmann, F. Stölzel, J.M. Middeke, M. Ciciarello, A. Curti, P.J.M. Valk, B. Löwenberg, M. Bornhäuser, J.F. DiPersio

### Analysis and interpretation of data

C. Lai, J. Vadakekolathu, S. Reeder, S.E. Church, T. Hood, G.A. Foulds, F. Stölzel, J.M. Middeke, P.J.M. Valk, B. Löwenberg, M. Bornhäuser, J.F. DiPersio, J.K. Davidson-Moncada, S. Rutella

### Clinical trial implementation

J.F. DiPersio was principal investigator at Washington University in St. Louis, St. Louis, United States of America. B. Löwenberg was principal investigator at Erasmus University Medical Centre, Rotterdam, Netherlands.

### Writing of the manuscript

C. Lai, S. Rutella

### Review and/or revision of the manuscript

C. Lai, J. Vadakekolathu, S. Reeder, S.E. Church, T. Hood, I. Aldoss, J. Godwin, M.J. Wieduwilt, M. Arellano, J. Muth, F. Ravandi, K. Sweet, H. Altmann, G.A. Foulds, F. Stölzel, J.M. Middeke, M. Ciciarello, A. Curti, P.J.M. Valk, B. Löwenberg, M. Bornhäuser, J.F. DiPersio, J.K. Davidson-Moncada, S. Rutella Study supervision: S. Rutella

## Data and materials availability

Processed input data and basic association analyses will be made available from the corresponding author on request for the purpose of conducting legitimate scientific research. The results shown in this paper are in part based upon data generated by the TCGA Research Network (https://www.cancer.gov/tcga).

## Supplemental Materials

**Supplemental Figure 1:**
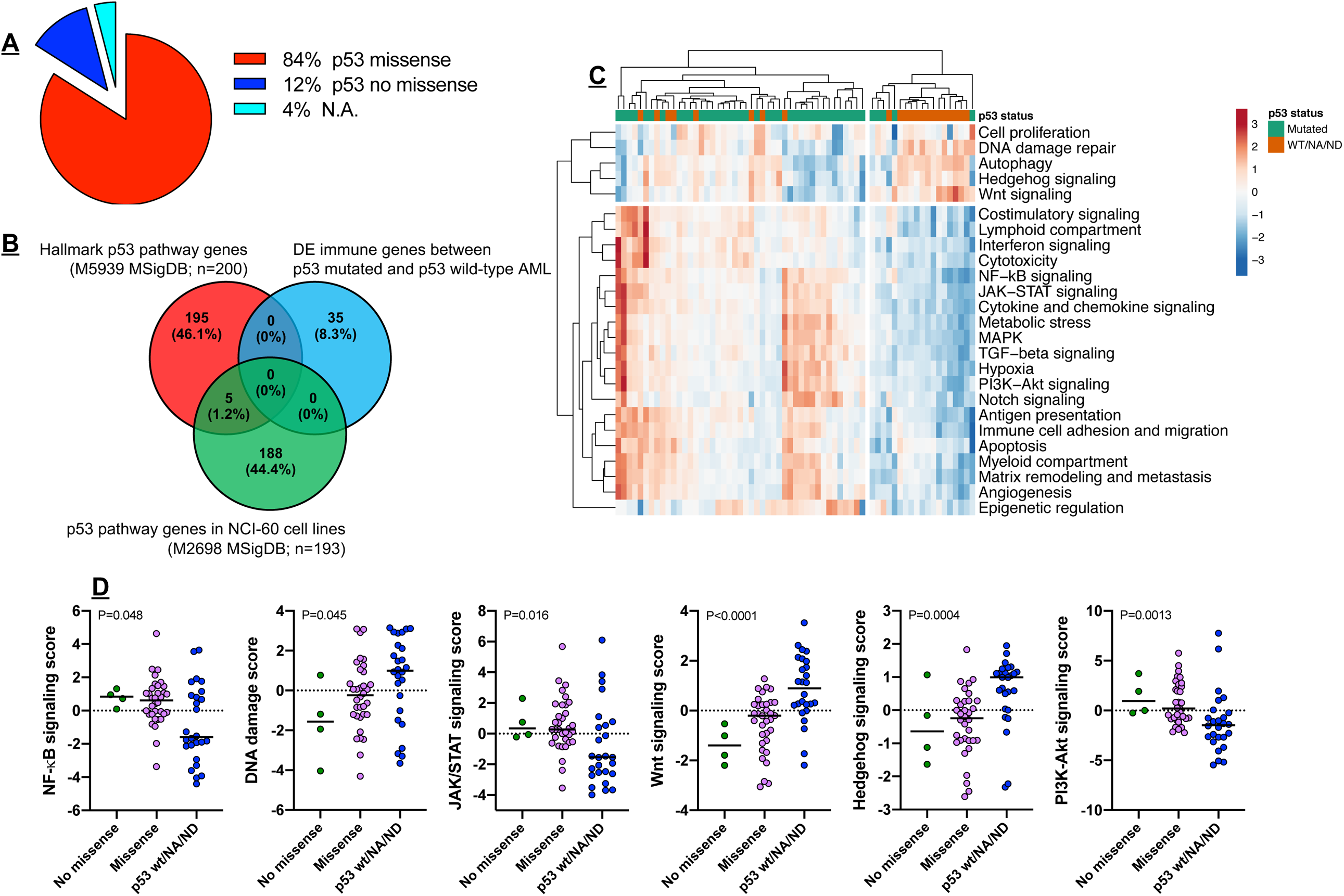
*TP53* mutational status and cancer pathways in the SAL cohort. **A**) *TP53* mutations were categorized as missense or no missense (frameshift, splice site and nonsense) using the IARC TP53 Database (http://p53.iarc.fr/) and based on prior knowledge (42,64). **B**) Venn diagram showing lack of overlap between differentially expressed (DE) immune genes in patients with *TP53* mutated AML and genes previously implicated in the *TP53* pathway. The Hallmark p53 Pathway Gene Set (n=200 genes involved in *TP53* pathways and networks; M5939) was downloaded from the Molecular Signature Database (MSigDB) (29,30). **C**) Heat-map of cancer pathway scores in patients with *TP53* mutations (n=42) and with *TP53* wild type AML (n=22; unsupervised hierarchical clustering; Euclidean distance; complete linkage). **D**) Cancer pathway scores in patients with *TP53* mutations and with *TP53* wild type AML. Bars denote median values. Data were compared using the Kruskal-Wallis test for unpaired determinations.

**Supplemental Figure 2:**
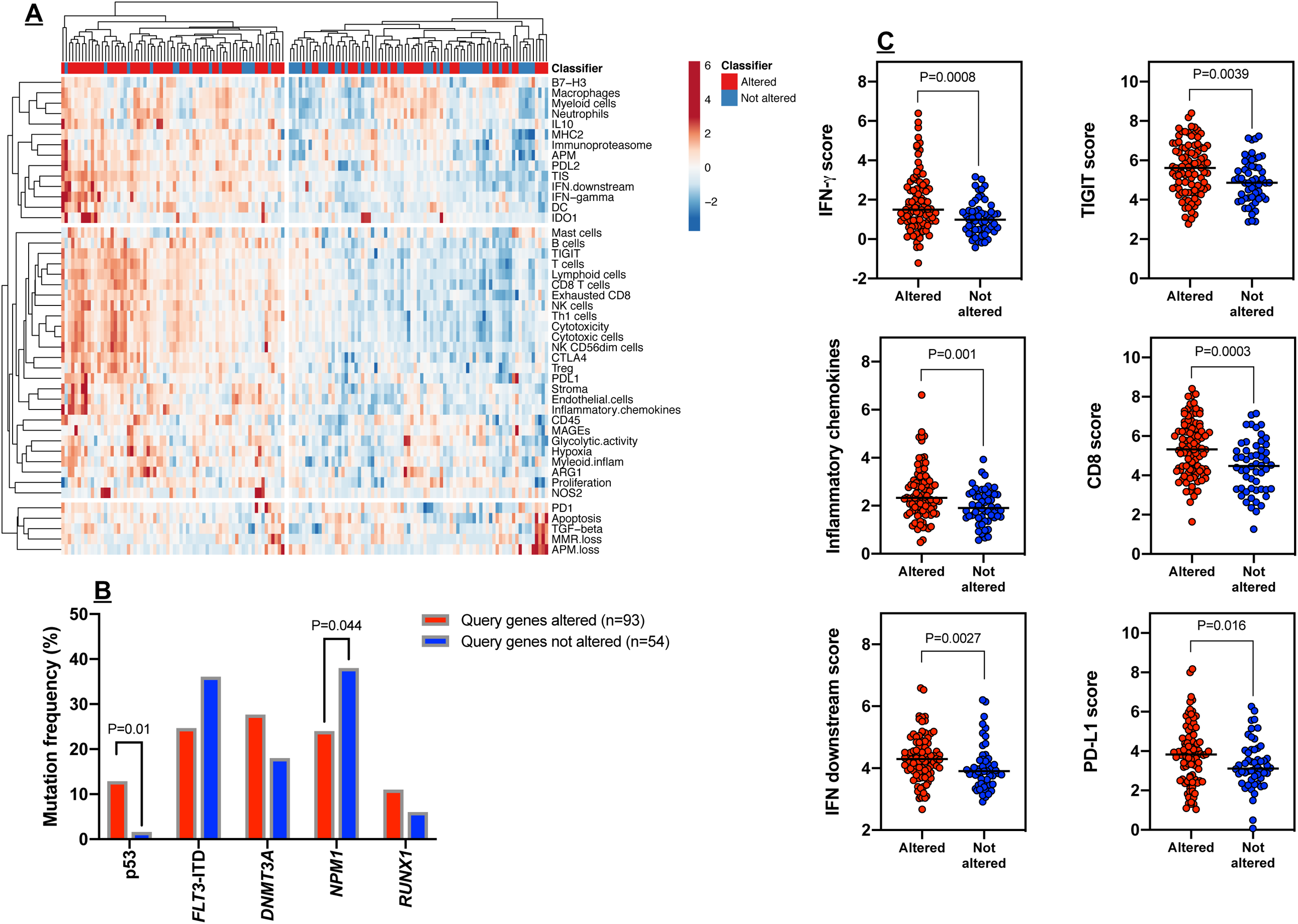
Genes in the *TP53* immune gene classifier correlate with immune infiltration in TCGA-AML cases. **A**) Heat-map of immune cell type-specific scores and biological activity scores (unsupervised hierarchical clustering; Euclidean distance; complete linkage) in TCGA-AML cases with (n=93) or without (n=54) abnormalities in the differentially expressed (DE) genes (*TP53* immune classifier) between patients with *TP53* mutated AML (n=42) and *TP53* wild type AML (n=22; **Fig. 2**). Abnormalities were defined as mRNA upregulation, gene amplification, deep deletion and mis-sense mutations relative to the gene’s expression distribution in all profiled AML samples. **B**) Correlation between abnormalities of the *TP53* immune gene classifier and prognostic molecular lesions, including *TP53* mutations, in TCGA-AML cases. Data were retrieved and analyzed using cBioPortal for Cancer Genomics (http://www.cbioportal.org/). **C**) Expression of IFN-γ signaling molecules, negative immune checkpoints and markers of T-cell infiltration in TCGA-AML cases with or without abnormalities in the *TP53* classifier genes. Bars denote median values. Data were compared using the Mann-Whitney *U* test for unpaired determinations.

**Supplemental Figure 3:**
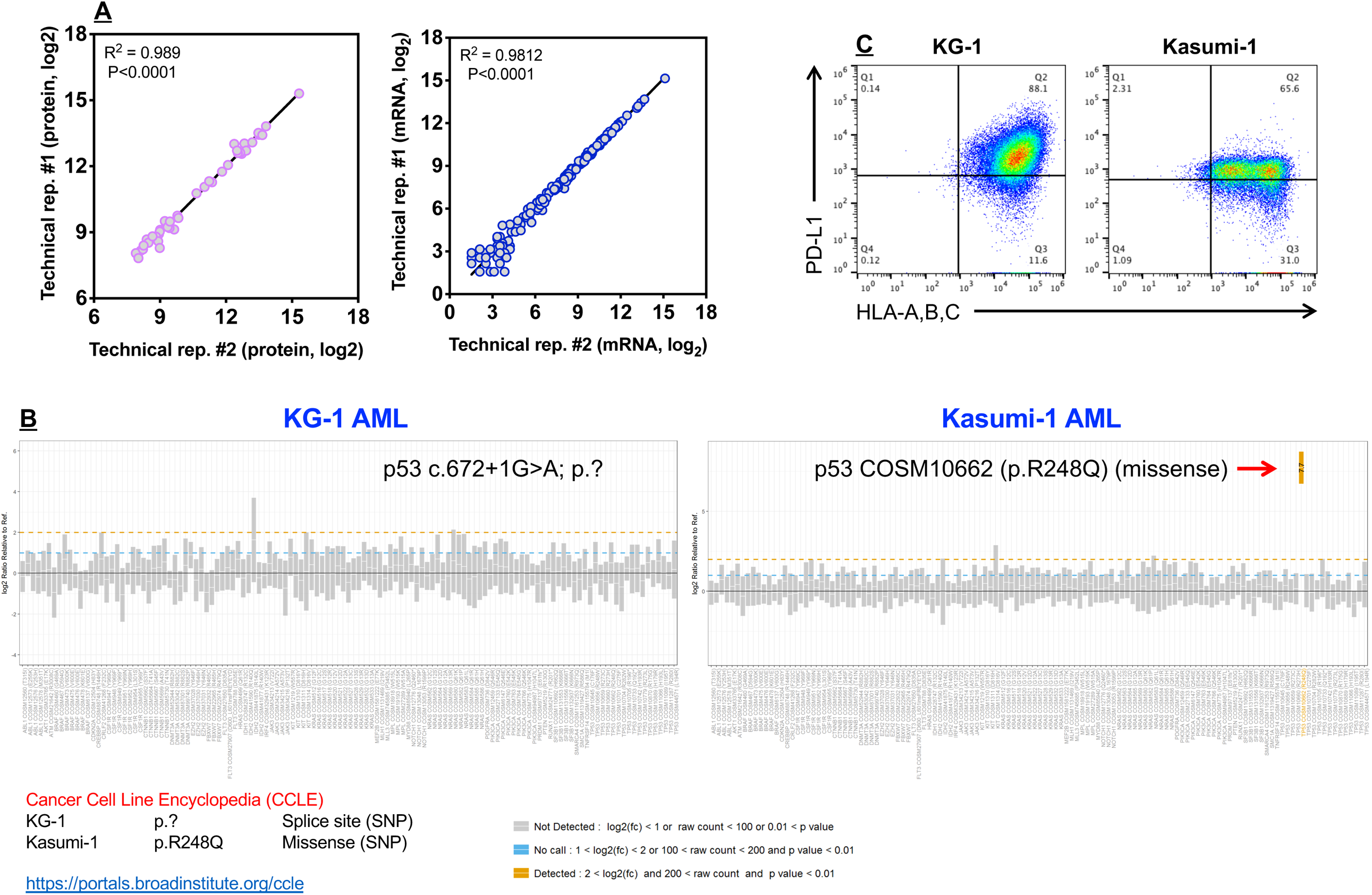
*TP53* status and expression of PD-L1 and MHC class I molecules in Kasumi-1 and KG-1 AML cell lines. **A**) Inter-assay reproducibility (replicate 1 and 2) of mRNA and protein measurements with the nCounter Vantage 3D™ Heme assay. R = Spearman correlation coefficient. **B**) Detection of single nucleotide variants (SNV) in Kasumi-1 and KG-1 cells using the nCounter Vantage 3D™ Heme assay (NanoString Technologies; for research use only and not for use in diagnostic procedures). **C**) Flow cytometric detection of PD-L1 and major histocompatibility complex (MHC) class I molecules in KG-1 and Kasumi-1 cells. Data were visualized using the FlowJo™ software package (version 10.6.1) and are representative of results in three independent experiments.

**Supplemental Table 1**: *TP53* mutations in TCGA-AML cases (n=147 patients).

**Supplemental Table 2**: *TP53* mutations in SAL-AML cases (n=40 patients).

**Supplemental Table 3**: Top differentially expressed genes (ranked by log_2_ fold change) between patients with *TP53* mutated (n=42) and *TP53* wild type AML (n=22).

**Supplemental Table 4**: Gene ontologies (GO) and KEGG pathways captured by differentially expressed (DE) genes between patients with *TP53* mutated (n=42) and *TP53* wild type AML (n=22).

**Supplemental Table 5**: Top differentially expressed genes/proteins (ranked by log_2_ fold change) between KG-1 (*TP53* loss-of-function [LOF]) and Kasumi-1 AML cells (*TP53* gain-of-function [GOF] mutation).

**Supplemental Table 6**: Gene ontologies (GO) and KEGG pathways captured by differentially expressed (DE) genes between KG-1 (*TP53* loss-of-function [LOF]) and Kasumi-1 AML (*TP53* gain-of-function [GOF]).

## References

1. Ravandi F, Cortes J, Faderl S, O’Brien S, Garcia-Manero G, Verstovsek S, et al. Characteristics and outcome of patients with acute myeloid leukemia refractory to 1 cycle of high-dose cytarabine-based induction chemotherapy. Blood 2010;116:5818-23; quiz 6153.

2. Hou HA, Chou WC, Kuo YY, Liu CY, Lin LI, Tseng MH, et al. TP53 mutations in de novo acute myeloid leukemia patients: longitudinal follow-ups show the mutation is stable during disease evolution. Blood Cancer J 2015;5:e331.

3. Hussaini MO, Mirza AS, Komrokji R, Lancet J, Padron E, Song J. Genetic landscape of acute myeloid leukemia interrogated by next-generation sequencing: A large cancer center experience. Cancer Genomics Proteomics 2018;15:121–6.

4. Seifert H, Mohr B, Thiede C, Oelschlagel U, Schakel U, Illmer T, et al. The prognostic impact of 17p (p53) deletion in 2272 adults with acute myeloid leukemia. Leukemia 2009;23:656–63.

5. Prokocimer M, Molchadsky A, Rotter V. Dysfunctional diversity of p53 proteins in adult acute myeloid leukemia: projections on diagnostic workup and therapy. Blood 2017;130:699–712.

6. Rucker FG, Schlenk RF, Bullinger L, Kayser S, Teleanu V, Kett H, et al. TP53 alterations in acute myeloid leukemia with complex karyotype correlate with specific copy number alterations, monosomal karyotype, and dismal outcome. Blood 2012;119:2114–21.

7. Stirewalt DL, Kopecky KJ, Meshinchi S, Appelbaum FR, Slovak ML, Willman CL, et al. FLT3, RAS, and TP53 mutations in elderly patients with acute myeloid leukemia. Blood 2001;97:3589–95.

8. Bowen D, Groves MJ, Burnett AK, Patel Y, Allen C, Green C, et al. TP53 gene mutation is frequent in patients with acute myeloid leukemia and complex karyotype, and is associated with very poor prognosis. Leukemia 2009;23:203–6.

9. Wei AH, Strickland SA, Jr., Hou JZ, Fiedler W, Lin TL, Walter RB, et al. Venetoclax combined with low-dose cytarabine for previously untreated patients with acute myeloid leukemia: Results from a phase Ib/II study. J Clin Oncol 2019;37:1277–84.

10. DiNardo CD, Pratz K, Pullarkat V, Jonas BA, Arellano M, Becker PS, et al. Venetoclax combined with decitabine or azacitidine in treatment-naive, elderly patients with acute myeloid leukemia. Blood 2019;133:7–17.

11. Ohgami RS, Ma L, Merker JD, Gotlib JR, Schrijver I, Zehnder JL, et al. Next-generation sequencing of acute myeloid leukemia identifies the significance of TP53, U2AF1, ASXL1, and TET2 mutations. Mod Pathol 2015;28:706–14.

12. Welch JS, Petti AA, Ley TJ. Decitabine in TP53-Mutated AML. N Engl J Med 2017;376:797–8.

13. Welch JS, Petti AA, Miller CA, Fronick CC, O’Laughlin M, Fulton RS, et al. TP53 and decitabine in acute myeloid leukemia and myelodysplastic syndromes. N Engl J Med 2016;375:2023–36.

14. Ciurea SO, Chilkulwar A, Saliba RM, Chen J, Rondon G, Patel KP, et al. Prognostic factors influencing survival after allogeneic transplantation for AML/MDS patients with TP53 mutations. Blood 2018;131:2989–92.

15. Munoz-Fontela C, Mandinova A, Aaronson SA, Lee SW. Emerging roles of p53 and other tumour-suppressor genes in immune regulation. Nat Rev Immunol 2016;16:741–50.

16. Zhang S, Zheng M, Kibe R, Huang Y, Marrero L, Warren S, et al. Trp53 negatively regulates autoimmunity via the STAT3-Th17 axis. FASEB J 2011;25:2387–98.

17. Komarova EA, Krivokrysenko V, Wang K, Neznanov N, Chernov MV, Komarov PG, et al. p53 is a suppressor of inflammatory response in mice. FASEB J 2005;19:1030–2.

18. Blagih J, Zani F, Chakravarty P, Hennequart M, Pilley S, Hobor S, et al. Cancer-specific loss of p53 leads to a modulation of myeloid and T cell responses. Cell Rep 2020;30:481–96 e6.

19. Hendrickx W, Simeone I, Anjum S, Mokrab Y, Bertucci F, Finetti P, et al. Identification of genetic determinants of breast cancer immune phenotypes by integrative genome-scale analysis. Oncoimmunology 2017;6:e1253654.

20. Cha YJ, Kim HR, Lee CY, Cho BC, Shim HS. Clinicopathological and prognostic significance of programmed cell death ligand-1 expression in lung adenocarcinoma and its relationship with p53 status. Lung Cancer 2016;97:73–80.

21. Wormann SM, Song L, Ai J, Diakopoulos KN, Kurkowski MU, Gorgulu K, et al. Loss of P53 function activates JAK2-STAT3 signaling to promote pancreatic tumor growth, stroma modification, and gemcitabine resistance in mice and is associated with patient survival. Gastroenterology 2016;151:180–93.

22. Dong ZY, Zhong WZ, Zhang XC, Su J, Xie Z, Liu SY, et al. Potential predictive value of TP53 and KRAS mutation status for response to PD-1 blockade immunotherapy in lung adenocarcinoma. Clin Cancer Res 2017;23:3012–24.

23. Rutella S, Church SE, Vadakekolathu J, Viboch E, Sullivan AH, Hood T, et al. Adaptive immune gene signatures correlate with response to flotetuzumab, a CD123 × CD3 bispecific DART® molecule, in patients with relapsed/refractory acute myeloid leukemia. Blood 2018;132:444-.

24. Boyiadzis M, Bishop MR, Abonour R, Anderson KC, Ansell SM, Avigan D, et al. The Society for Immunotherapy of Cancer consensus statement on immunotherapy for the treatment of hematologic malignancies: multiple myeloma, lymphoma, and acute leukemia. J Immunother Cancer 2016;4:90.

25. Ley TJ, Miller C, Ding L, Raphael BJ, Mungall AJ, Robertson A, et al. Genomic and epigenomic landscapes of adult de novo acute myeloid leukemia. N Engl J Med 2013;368:2059–74.

26. Valk PJ, Verhaak RG, Beijen MA, Erpelinck CA, Barjesteh van Waalwijk van Doorn-Khosrovani S, Boer JM, et al. Prognostically useful gene-expression profiles in acute myeloid leukemia. N Engl J Med 2004;350:1617–28.

27. Stavropoulou V, Kaspar S, Brault L, Sanders MA, Juge S, Morettini S, et al. MLL-AF9 expression in hematopoietic stem cells drives a highly invasive AML expressing EMT-related genes linked to poor outcome. Cancer Cell 2016;30:43–58.

28. Wagner S, Vadakekolathu J, Tasian SK, Altmann H, Bornhauser M, Pockley AG, et al. A parsimonious 3-gene signature predicts clinical outcomes in an acute myeloid leukemia multicohort study. Blood Adv 2019;3:1330–46.

29. Danaher P, Warren S, Cesano A. Development of gene expression signatures characterizing the tumor-immune interaction. J Clin Oncol 2018;36:205-.

30. Danaher P, Warren S, Dennis L, D’Amico L, White A, Disis ML, et al. Gene expression markers of Tumor Infiltrating Leukocytes. J Immunother Cancer 2017;5:18.

31. Subramanian A, Tamayo P, Mootha VK, Mukherjee S, Ebert BL, Gillette MA, et al. Gene set enrichment analysis: a knowledge-based approach for interpreting genome-wide expression profiles. Proc Natl Acad Sci U S A 2005;102:15545–50.

32. Bezzi M, Seitzer N, Ishikawa T, Reschke M, Chen M, Wang G, et al. Diverse genetic-driven immune landscapes dictate tumor progression through distinct mechanisms. Nat Med 2018;24:165–75.

33. Versluis J, In ‘t Hout FE, Devillier R, van Putten WL, Manz MG, Vekemans MC, et al. Comparative value of post-remission treatment in cytogenetically normal AML subclassified by NPM1 and FLT3-ITD allelic ratio. Leukemia 2017;31:26–33.

34. Ott PA, Bang YJ, Piha-Paul SA, Razak ARA, Bennouna J, Soria JC, et al. T-cell-inflamed gene-expression profile, programmed death ligand 1 expression, and tumor mutational burden predict efficacy in patients treated with pembrolizumab across 20 cancers: KEYNOTE-028. J Clin Oncol 2019;37:318–27.

35. Ayers M, Lunceford J, Nebozhyn M, Murphy E, Loboda A, Kaufman DR, et al. IFN-gamma-related mRNA profile predicts clinical response to PD-1 blockade. J Clin Invest 2017;127:2930–40.

36. Bouaoun L, Sonkin D, Ardin M, Hollstein M, Byrnes G, Zavadil J, et al. TP53 variations in human cancers: New lessons from the IARC TP53 database and genomics data. Hum Mutat 2016;37:865–76.

37. Georgoudaki AM, Prokopec KE, Boura VF, Hellqvist E, Sohn S, Ostling J, et al. Reprogramming tumor-associated macrophages by antibody targeting inhibits cancer progression and metastasis. Cell Rep 2016;15:2000–11.

38. La Fleur L, Boura VF, Alexeyenko A, Berglund A, Ponten V, Mattsson JSM, et al. Expression of scavenger receptor MARCO defines a targetable tumor-associated macrophage subset in non-small cell lung cancer. Int J Cancer 2018;143:1741–52.

39. Knaus HA, Berglund S, Hackl H, Blackford AL, Zeidner JF, Montiel-Esparza R, et al. Signatures of CD8+ T cell dysfunction in AML patients and their reversibility with response to chemotherapy. JCI Insight 2018;3:e120974.

40. Kim NH, Kim HS, Kim NG, Lee I, Choi HS, Li XY, et al. p53 and microRNA-34 are suppressors of canonical Wnt signaling. Sci Signal 2011;4:ra71.

41. Boettcher S, Miller PG, Sharma R, McConkey M, Leventhal M, Krivtsov AV, et al. A dominant-negative effect drives selection of TP53 missense mutations in myeloid malignancies. Science 2019;365:599–604.

42. Schulz-Heddergott R, Stark N, Edmunds SJ, Li J, Conradi LC, Bohnenberger H, et al. Therapeutic ablation of gain-of-function mutant p53 in colorectal cancer inhibits Stat3-mediated tumor growth and invasion. Cancer Cell 2018;34:298–314.

43. Vadakekolathu J, Minden MD, Hood T, Church SE, Reeder S, Altmann H, et al. Immune landscapes predict chemotherapy resistance and anti-leukemic activity of flotetuzumab, an investigational CD123×CD3 bispecific Dart® molecule, in patients with relapsed/refractory acute myeloid leukemia. Blood 2019;134:460-.

44. Chichili GR, Huang L, Li H, Burke S, He L, Tang Q, et al. A CD3xCD123 bispecific DART for redirecting host T cells to myelogenous leukemia: preclinical activity and safety in nonhuman primates. Sci Transl Med 2015;7:289ra82.

45. Miller C, Mohandas T, Wolf D, Prokocimer M, Rotter V, Koeffler HP. Human p53 gene localized to short arm of chromosome 17. Nature 1986;319:783–4.

46. Miller LD, Smeds J, George J, Vega VB, Vergara L, Ploner A, et al. An expression signature for p53 status in human breast cancer predicts mutation status, transcriptional effects, and patient survival. Proc Natl Acad Sci U S A 2005;102:13550–5.

47. Munoz-Fontela C, Pazos M, Delgado I, Murk W, Mungamuri SK, Lee SW, et al. p53 serves as a host antiviral factor that enhances innate and adaptive immune responses to influenza A virus. J Immunol 2011;187:6428–36.

48. Cooks T, Pateras IS, Tarcic O, Solomon H, Schetter AJ, Wilder S, et al. Mutant p53 prolongs NF-kappaB activation and promotes chronic inflammation and inflammation-associated colorectal cancer. Cancer Cell 2013;23:634–46.

49. Cooks T, Pateras IS, Jenkins LM, Patel KM, Robles AI, Morris J, et al. Mutant p53 cancers reprogram macrophages to tumor supporting macrophages via exosomal miR-1246. Nat Commun 2018;9:771.

50. Quigley D, Silwal-Pandit L, Dannenfelser R, Langerod A, Vollan HK, Vaske C, et al. Lymphocyte invasion in IC10/basal-like breast tumors is associated with wild-type TP53. Mol Cancer Res 2015;13:493–501.

51. Liu Z, Jiang Z, Gao Y, Wang L, Chen C, Wang X. TP53 mutations promote immunogenic activity in breast cancer. J Oncol 2019;2019:5952836.

52. Brosh R, Rotter V. When mutants gain new powers: news from the mutant p53 field. Nat Rev Cancer 2009;9:701–13.

53. Hayashi Y, Goyama S, Liu X, Tamura M, Asada S, Tanaka Y, et al. Antitumor immunity augments the therapeutic effects of p53 activation on acute myeloid leukemia. Nat Commun 2019;10:4869.

54. Iannello A, Thompson TW, Ardolino M, Lowe SW, Raulet DH. p53-dependent chemokine production by senescent tumor cells supports NKG2D-dependent tumor elimination by natural killer cells. J Exp Med 2013;210:2057–69.

55. Malekzadeh P, Pasetto A, Robbins PF, Parkhurst MR, Paria BC, Jia L, et al. Neoantigen screening identifies broad TP53 mutant immunogenicity in patients with epithelial cancers. J Clin Invest 2019;129:1109–14.

56. Malekzadeh P, Yossef R, Cafri G, Paria BC, Lowery FJ, Jafferji M, et al. Antigen experienced T cells from peripheral blood recognize p53 neoantigens. Clin Cancer Res 2020.

57. Ham SW, Jeon HY, Jin X, Kim EJ, Kim JK, Shin YJ, et al. TP53 gain-of-function mutation promotes inflammation in glioblastoma. Cell Death Differ 2019;26:409–25.

58. Horibata S, Gui G, Lack J, DeStefano CB, Gottesman MM, Hourigan CS. Heterogeneity in refractory acute myeloid leukemia. Proc Natl Acad Sci U S A 2019;116:10494–503.

59. Quintas-Cardama A, Hu C, Qutub A, Qiu YH, Zhang X, Post SM, et al. p53 pathway dysfunction is highly prevalent in acute myeloid leukemia independent of TP53 mutational status. Leukemia 2017;31:1296–305.

60. Ley TJ, Ding L, Walter MJ, McLellan MD, Lamprecht T, Larson DE, et al. DNMT3A mutations in acute myeloid leukemia. N Engl J Med 2010;363:2424–33.

61. Ghoneim HE, Fan Y, Moustaki A, Abdelsamed HA, Dash P, Dogra P, et al. De novo epigenetic programs inhibit PD-1 blockade-mediated T cell rejuvenation. Cell 2017;170:142–57 e19.

62. Mathew NR, Baumgartner F, Braun L, O’Sullivan D, Thomas S, Waterhouse M, et al. Sorafenib promotes graft-versus-leukemia activity in mice and humans through IL-15 production in FLT3-ITD-mutant leukemia cells. Nat Med 2018;24:282–91.

63. Metsalu T, Vilo J. ClustVis: a web tool for visualizing clustering of multivariate data using Principal Component Analysis and heatmap. Nucleic Acids Res 2015;43:W566–70.

64. Muller PA, Vousden KH. Mutant p53 in cancer: new functions and therapeutic opportunities. Cancer Cell 2014;25:304–17.

65. Tang Z, Kang B, Li C, Chen T, Zhang Z. GEPIA2: an enhanced web server for large-scale expression profiling and interactive analysis. Nucleic Acids Res 2019;47:W556–W60.

