## Supplemental Materials for "*TP53* abnormalities correlate with immune infiltration and are associated with response to flotetuzumab, an investigational immunotherapy, in acute myeloid leukemia"

| **Sample ID** | **Protein change** | **Mutation type*** |
| --- | --- | --- |
| TCGA-AB-2943-03 | R273C | Missense |
| TCGA-AB-2935-03 | R248Q | Missense |
| TCGA-AB-2813-03 | C176Y | Missense |
| TCGA-AB-2938-03 | H179R | Missense |
| TCGA-AB-2908-03 | C141W | Missense |
| TCGA-AB-2908-03 | Q317* | Nonsense |
| TCGA-AB-2885-03 | H193Y | Missense |
| TCGA-AB-2829-03 | R280G | Missense |
| TCGA-AB-2829-03 | X225_splice | Splice site |
| TCGA-AB-2904-03 | R337C | Missense |
| TCGA-AB-2952-03 | E286G | Missense |
| TCGA-AB-2941-03 | I195S | Missense |
| TCGA-AB-2878-03 | V173* | Frameshift |
| TCGA-AB-2878-03 | S215G | Missense |
| TCGA-AB-2820-03 | P223Rfs*4 | Frameshift |
| TCGA-AB-2860-03 | M40Lfs*7 | Frameshift |
| TCGA-AB-2838-03 | T125= | Splice site |
| TCGA-AB-2938-03 | R342Efs*3 | Frameshift |
| TCGA-AB-2868-03 | X126_splice | Splice site |
| TCGA-AB-2857-03 | X10_splice | Splice site |

**Supplemental Table 1**: Somatic *TP53* mutations in TCGA-AML cases (n=147 patients).

*IARC *TP53* Database. Two different *TP53* mutations were identified in 4 patients from this series (green highlight).

**Supplemental Table 2**: Somatic *TP53* mutations in SAL-AML cases (n=40 patients).

| **Subject ID** | **Protein change** | **Mutation type*** |
| --- | --- | --- |
| 32-2-001 | G360E | N.A. |
| 02-1-053 | R181C | Missense |
| 03-1-006 | T211I | Missense |
| 03-1-120 | P151S | Missense |
| 03-1-166 | H178P | Missense |
| 03-1-174 | [STOP] AA | Frameshift |
| 03-1-225 | T125M | Missense |
| 08-1-024 | R175H | Missense |
| 09-1-051 | K132T | Missense |
| 09-1-077 | C238Y | Missense |
| 09-1-114 | V272M | Missense |
| 09-1-130 | R175G | Missense |
| 09-1-143 | R248Q | Missense |
| 09-1-222 | R248Q | Missense |
| 10-1-012 | E286K | Missense |
| 10-1-024 | R248W | Missense |
| 10-1-082 | I195T | Missense |
| 12-1-002 | H179Y | Missense |
| 15-1-018 | [STOP] AA 344 | Frameshift |
| 15-1-025 | [STOP] AA 125 | Frameshift |
| 16-1-031 | R280G | Missense |
| 23-1-026 | I195N | Missense |
| 28-1-020 | P278R | Missense |
| 29-1-010 | M237K | Missense |
| 30-1-008 | M246V | Missense |
| 30-1-008 | R283C | Missense |
| 30-1-130 | Y ® [STOP] | Frameshift |
| 33-1-003 | H193R | Missense |
| 35-1-022 | F270C | Missense |
| 37-1-006 | R283C | Missense |
| 39-1-027 | [STOP] AA 169 | Frameshift |
| 39-1-027 | Y220C | Missense |
| 007-001-115 | R283QR | N.A. |
| 036-006-154 | N235S | Missense |
| 046-021-326 | V216M | Missense |
| R-S-014-154-00RH | P151T | Missense |
| R-S-030-140-CC3U | R273H | Missense |
| R-S-039-025-ECR7 | E286G | Missense |
| R-S-052-018-N3CL | V272M | Missense |
| R-S-054-137-44VL | G245V | Missense |
| R-S-067-024-VSLZ | N235S | Missense |
| R-S-030-176-ZZQH | C275Y | Missense |

*IARC *TP53* Database. Two different *TP53* mutations were identified in 2 patients from this series (green highlight).

**Supplemental Table 3**: Top differentially expressed genes (ranked by log_2_ fold change) between patients with *TP53* mutated and *TP53* wild type AML. Results were filtered to include only those genes that displayed ≥1.5-fold changes in expression and had passed a test for statistical significance (P<0.01).

| **Gene Id** | **Log2 fold change** | **P value** |
| --- | --- | --- |
| ***Up in TP53 mutant*** | | |
| THBD | 3.28 | 2.88×10^-10^ |
| IL33 | 2.67 | 4.31×10^-6^ |
| CCL3/L1 | 2.61 | 1.44×10^-6^ |
| IL6 | 2.32 | 0.0001396 |
| CXCL8 | 2.25 | 2.27×10^-5^ |
| RIPK2 | 2.15 | 9.04×10^-13^ |
| CCL2 | 2.12 | 0.0283116 |
| THBS1 | 2.10 | 0.0061577 |
| CXCL1 | 2.09 | 0.0001883 |
| CSF1 | 2.01 | 2.65×10^-5^ |
| MYCT1 | 1.97 | 9.35×10^-6^ |
| OASL | 1.92 | 0.0002112 |
| PTGER4 | 1.92 | 2.37×10^-10^ |
| CXCL2 | 1.82 | 2.65×10^-5^ |
| BBC3 | 1.81 | 1.64×10^-6^ |
| DUSP5 | 1.77 | 1.48×10^-7^ |
| IFNG | 1.73 | 0.000511 |
| FCAR | 1.71 | 0.001586 |
| ***Up in TP53 wild type*** | | |
| CD2 | -1.61 | 3.91×10^-8^ |
| CD79A | -1.73 | 0.0003015 |
| CD79B | -1.76 | 0.0006327 |
| GIMAP6 | -1.88 | 6.53×10^-5^ |
| CCR2 | -1.89 | 3.28×10^-5^ |
| PRF1 | -1.94 | 0.0003822 |
| FAM30A | -1.97 | 0.0250546 |
| NT5E | -2.01 | 9.74×10^-7^ |
| ITGB3 | -2.09 | 1.07×10^-6^ |
| CD19 | -2.20 | 1.49×10^-5^ |
| CXCR2 | -2.32 | 2.92×10^-5^ |
| MS4A1 | -3.40 | 2.25×10^-9^ |
| GBP1 | -3.57 | 1.38×10^-6^ |
| VTCN1 | -3.88 | 6.96×10^-22^ |
| MARCO | -3.96 | 2.30×10^-7^ |
| CX3CR1 | -4.72 | 2.24×10^-8^ |

**Supplemental Table 4**: Gene ontologies (GO) and KEGG pathways captured by differentially expressed (DE) genes between patients with *TP53* mutated and *TP53* wild type AML. FDR = false discovery rate.

| **GO term** | **Description** | **Count in gene set** | **FDR** |
| --- | --- | --- | --- |
| GO:0006955 | Immune response | 21 of 1560 | 2.94×10^-15^ |
| GO:0032101 | Regulation of response to external stimulus | 17 of 732 | 3.21×10^-15^ |
| GO:0071345 | Cellular response to cytokine stimulus | 18 of 953 | 4.67×10^-15^ |
| GO:0006952 | Defense response | 19 of 1234 | 9.97×10^-15^ |
| GO:0032103 | Positive regulation of response to external stimulus | 13 of 291 | 1.18×10^-14^ |
| GO:0002376 | Immune system process | 22 of 2370 | 7.55×10^-14^ |
| GO:0006950 | Response to stress | 24 of 3267 | 8.40×10^-14^ |
| GO:0006954 | Inflammatory response | 14 of 482 | 8.99×10^-14^ |
| GO:0019221 | Cytokine-mediated signaling pathway | 15 of 655 | 1.55×10^-13^ |
| **Pathway** | **Description** | **Count in gene set** | **FDR** |
| hsa04060 | Cytokine-cytokine receptor interaction | 9 of 263 | 5.56×10^-9^ |
| hsa05323 | Rheumatoid arthritis | 6 of 84 | 6.37×10^-8^ |
| hsa04657 | IL-17 signaling pathway | 6 of 92 | 8.02×10^-8^ |
| hsa04621 | NOD-like receptor signaling pathway | 6 of 166 | 1.56×10^-6^ |
| hsa04668 | TNF signaling pathway | 5 of 108 | 4.16×10^-6^ |
| hsa05200 | Pathways in cancer | 7 of 515 | 4.09×10^-5^ |
| hsa04151 | PI3K-Akt signaling pathway | 5 of 348 | 0.00056 |
| hsa04062 | Chemokine signaling pathway | 4 of 181 | 0.00056 |
| hsa04659 | Th17 cell differentiation | 3 of 102 | 0.0017 |

**Supplemental Table 5**: Top differentially expressed genes/proteins (ranked by log_2_ fold change) between KG-1 (*TP53* loss-of-function [LOF]) and Kasumi-1 AML cells (*TP53* gain-of-function [GOF] mutation). Results were filtered to include only those genes that displayed ≥1.7-fold changes in expression and had passed a test for statistical significance (P<0.01).

| **Gene Id** | **Log2 fold change** | **P value** |
| --- | --- | --- |
| ***Up in Kasumi-1 cells*** | | |
| p53 protein | 10.0 | 0.00029 |
| FYN-mRNA | 5.56 | 0.00084 |
| DNMT3A-mRNA | 2.35 | 5.67×10^-5^ |
| CSF3R-mRNA | 2.32 | 4.43×10^-5^ |
| DDIT3-mRNA | 2.16 | 0.00085 |
| KIT-mRNA | 2.08 | 0.00014 |
| CBL-mRNA | 1.89 | 3.16×10^-6^ |
| FOXO1-mRNA | 1.83 | 0.0051 |
| TP53-mRNA | 1.73 | 0.0079 |
| STAT6-mRNA | 1.70 | 8.54×10^-6^ |
| CDKN2A-mRNA | 1.70 | 0.00021 |
| ***Up in KG-1 cells*** | | |
| EIF4E-mRNA | -1.81 | 0.00029 |
| IFI16-mRNA | -1.91 | 1.42×10^-5^ |
| JAK2-mRNA | -1.91 | 0.015 |
| CHEK1-mRNA | -1.96 | 0.0071 |
| NLRC5-mRNA | -1.99 | 0.00027 |
| Iκ-Ba-protein | -2.06 | 0.00031 |
| CD79A-mRNA | -2.08 | 0.0143 |
| KAT6A-mRNA | -2.13 | 1.68×10^-5^ |
| IRF1-mRNA | -2.19 | 2.18×10^-5^ |
| RHOA-mRNA | -2.21 | 3.94×10^-5^ |
| IKKβ-protein | -2.24 | 3.17×10^-6^ |
| IDH1-mRNA | -2.36 | 8.95×10^-5^ |
| BCL3-mRNA | -2.50 | 0.00034 |
| STAT1-mRNA | -2.57 | 1.41×10^-5^ |
| RARA-mRNA | -2.64 | 0.00041 |
| CCND3-mRNA | -2.74 | 9.71×10^-5^ |
| STAT5A-mRNA | -3.0 | 1.86×10^-5^ |
| CCND2-mRNA | -3.1 | 6.35×10^-5^ |
| SYK-mRNA | -3.48 | 2.84×10^-6^ |
| IL2RA-mRNA | -3.63 | 0.017 |
| RAC1-mRNA | -3.75 | 4.02×10^-6^ |
| LYN-mRNA | -3.78 | 1.32×10^-6^ |
| LCK-mRNA | -4.02 | 0.00285 |
| OSM-mRNA | -4.38 | 0.0059 |
| CD34-mRNA | -4.39 | 4.43×10^-7^ |
| SET-mRNA | -4.71 | 7.37×10^-6^ |
| MYCN-mRNA | -4.76 | 0.00302 |
| PIM1-mRNA | -4.95 | 2.65×10^-6^ |
| HOXA9-mRNA | -5.24 | 0.00014 |
| NF-κB-protein | -5.33 | 0.0028 |
| IKZF2-mRNA | -5.79 | 2.46×10^-5^ |
| MEIS1-mRNA | -6.99 | 0.0034 |
| CIITA-mRNA | -7.10 | 0.00133 |
| HGF-mRNA | -7.25 | 0.00118 |

**Supplemental Table 6**: Gene ontologies (GO) and KEGG pathways captured by differentially expressed (DE) genes between KG-1 (*TP53* loss-of-function [LOF]) and Kasumi-1 AML (*TP53* gain-of-function [GOF]). FDR = false discovery rate.

| **GO term** | **Description** | **Count in gene set** | **FDR** |
| --- | --- | --- | --- |
| GO:0010604 | Positive regulation of macromolecule metabolic process | 33 of 3081 | 1.17×10^-14^ |
| GO:0005515 | Protein binding | 41 of 6605 | 5.58×10^-14^ |
| GO:0004672 | Protein kinase activity | 15 of 635 | 7.21×10^-10^ |
| GO:0016301 | Kinase activity | 16 of 835 | 1.71×10^-9^ |
| GO:0010604 | Positive regulation of macromolecule metabolic process | 33 of 3081 | 1.17×10^-14^ |
| GO:0051171 | Regulation of nitrogen compound metabolic process | 39 of 5827 | 1.39×10^-13^ |
| GO:0051171 | Regulation of nitrogen compound metabolic process | 39 of 5827 | 1.39×10^-13^ |
| GO:0002682 | Regulation of immune system process | 21 of 1391 | 3.36×10^-11^ |
| GO:0016310 | Phosphorylation | 20 of 1236 | 4.28×10^-11^ |
| **Pathway** | **Description** | **Count in gene set** | **FDR** |
| hsa05203 | Viral carcinogenesis | 15 of 183 | 1.16×10^-17^ |
| hsa05200 | Pathways in cancer | 18 of 515 | 1.80×10^-15^ |
| hsa04151 | PI3K-Akt signaling pathway | 15 of 348 | 4.93×10^-14^ |
| hsa04218 | Cellular senescence | 10 of 156 | 4.21×10^-11^ |
| hsa04630 | JAK-STAT signaling pathway | 10 of 160 | 4.81×10^-11^ |
| hsa04659 | Th17 cell differentiation | 8 of 102 | 1.10×10^-9^ |
| hsa05221 | Acute myeloid leukemia | 7 of 66 | 2.30×10^-9^ |
| hsa04115 | p53 signaling pathway | 7 of 68 | 2.61×10^-9^ |
| hsa04658 | Th1 and Th2 cell differentiation | 7 of 88 | 1.33×10^-8^ |
